# Long-term ketogenic diet causes hyperlipidemia, liver dysfunction, and glucose intolerance from impaired insulin trafficking and secretion in mice

**DOI:** 10.1101/2024.06.14.599117

**Authors:** Molly R. Gallop, Renan F.L. Vieira, Elijah T. Matsuzaki, Peyton D. Mower, Willisa Liou, Faith E. Smart, Seth Roberts, Kimberley J. Evason, William L. Holland, Amandine Chaix

## Abstract

A ketogenic diet (KD) is a very low-carbohydrate, very high-fat diet proposed to treat obesity and type 2 diabetes. While KD grows in popularity, its effects on metabolic health are understudied. Here we show that, in male and female mice, while KD protects against weight gain and induces weight loss, over long-term, mice develop hyperlipidemia, hepatic steatosis, and severe glucose intolerance. Unlike high fat diet-fed mice, KD mice are not insulin resistant and have low levels of insulin. Hyperglycemic clamp and *ex vivo* GSIS revealed cell-autonomous and whole-body impairments in insulin secretion. Major ER/Golgi stress and disrupted ER-Golgi protein trafficking was indicated by transcriptomic profiling of KD islets and confirmed by electron micrographs showing a dilated Golgi network likely responsible for impaired insulin granule trafficking and secretion. Overall, our results suggest long-term KD leads to multiple aberrations of metabolic parameters that caution its systematic use as a health promoting dietary intervention.

## Introduction

A ketogenic diet (KD) is a very high fat low carbohydrate diet that has been used to manage refractory epilepsy for the past hundred years.^1,2^ The exact mechanisms by which a KD confers seizure control are unknown, but there are a few hypotheses including reduction in blood glucose levels, favorable changes to brain metabolism, neurotransmitter levels and synaptic activity, and anti-convulsant effects of ketone bodies themselves.^3-5^ A KD does not have a defined macronutrient content, but, as a treatment for epilepsy, it is generally first prescribed at a 4:1 ratio of 4 grams of fat for every gram of carbohydrate or protein translating to roughly 90% of calories coming from fat.^2,6^ This high proportion of energy intake from fat causes the body to use fat rather than glucose as the primary fuel source and to enter a state of increased ketone body production, known as ketosis.^1,7^

Ketone bodies are produced primarily in the liver when glucose and insulin levels are low such as during prolonged fasting or when consuming a ketogenic diet. In the 1960s, it was proposed that this state of low glucose and insulin associated with KD consumption could treat obesity and related metabolic health conditions by favoring fat usage over fat storage.^7^ In addition, for similar insulin and glucose lowering reasons as well a potential benefits of ketone bodies themselves, a KD has been proposed to treat cancer, Alzheimer’s disease, increase longevity, and improve metabolic health.^8,9^ However, a breadth of evidence to support these claims is lacking. A recent study found that a KD increases cellular markers of aging in mice,^10^ and in 2019 JAMA released a publication titled “The Ketogenic Diet for Obesity and Diabetes— Enthusiasm Outpaces Evidence.”^11^ While a KD helps to control seizures in some people, its benefits will not necessarily translate to treating other diseases or disorders. In particular, pre-existing conditions, such as obesity and diabetes, might interact with a KD in ways that have not been fully characterized in the predominantly young, and limited group of epilepsy patients that have typically been studied.^6,12^ Therefore, harmful interactions between a KD and metabolic health conditions may negate the generally accepted belief that a KD is a clinically safe dietary intervention. From a nutritional and cardiometabolic standpoint, it is conceivable that there could be long-term ramifications to consuming a diet primarily composed of fat with very few carbohydrates that have been largely underexplored.^13,14^

Previous studies in mice fed a KD have reported conflicting findings on metabolic health. For example, compared to mice on a chow diet, most studies observed a lower body weight on KD ^15-19^ but others found that a KD caused weight gain.^9,20^ In contrast to the weight benefits found in many KD studies, high levels of blood cholesterol and non-esterified fatty acids (NEFA) have been observed in multiple strains of mice on KD.^15,17,19,21,22^ Furthermore, the effects of KD on glucose homeostasis have led to discordant and partial results. While some studies reported improvements in static blood glucose without a glucose challenge^18,21,23^ and improved glucose tolerance in WT but also obese and diabetic models (ob, db, stz)^16,24^, others reported insulin resistance,^25^ glucose intolerance,^26,27^ and loss of β cells.^26,27^ Importantly, the lack of consistency between dietary macronutrient composition of KDs and differences in food intake could potentially explain incongruencies. Overall, possible CVD risk associated with heightened lipid levels and inconclusive results on glucose homeostasis warrant further investigations into the long-term effect of KD consumption to define the conditions in which it can be helpful or harmful.

Another important factor when considering the effects of KD resides in the study design, namely whether KD was used to prevent versus treat metabolic health conditions. In humans, KD is a very popular weight loss (WL) intervention and has also been show to induce weight loss in mice.^15,28,29^ However, WL per se may not be sufficient to normalize metabolic complications. For example, serum dyslipidemia and liver steatosis seem to persist despite reductions in weight under KD. Moreover, the effects of KD-induced WL on glucose regulation are also unclear: some studies noted improvements in glucose tolerance testing^15,28,29^ while some did not.^30^ Noticeably, hypoglycemia, a known complication of treating type 2 diabetes, was observed in mice on KD,^24,28,29^ which raises concerns about the safety of using KD in a population taking diabetes medication.^31^ Finally, studies of KD-induced WL describe a reduction in HFD-linked hyperinsulinemia, yet we noted that in most, insulin levels dropped below that of chow-fed animals, which could be detrimental in the context of regulating fat storage and gluconeogenesis. Our search for preclinical studies characterizing KD-mediated WL only identified five studies thus far.^15,24,28-30^ With the mainstream popularity of KD as a therapeutic for obesity and diabetes, it is critical to further establish the broad metabolic effects of WL under KD, its effects on pre-existing metabolic conditions, as well as possible sex differences in these responses using a well-defined diet and a systematic metabolic characterization.

This study evaluated the effects of a long-term ketosis-causing KD on overall metabolic health in male and female C57Bl/6J mice to evaluate sex-differences in the response to KD feeding. The effect of a KD on body weight and on multiple parameters of metabolic health was compared to that of standard 60% HFD, a standard low-fat diet, and a moderate protein diet that protein matched to the KD. We also tested the metabolic effect of KD-induced WL in obese males and females.

## Results

### 1 A 90% fat KD prevents weight gain as compared to a HFD in both male and female mice

Adult C57Bl/6J male or female mice were placed on one of four long-term dietary interventions– a 10% fat low fat diet (LFD), a standard 60% high fat diet (HFD), an 89.9% fat ketogenic diet, or another 10% fat low-fat diet with 10% protein instead of the standard 20% to match the protein content of the ketogenic diet (low fat moderate protein-LFMP) (**Figure 1A**,**1B**). To test whether our KD induced ketosis, we compared fasted and fed levels of blood ketone body β-hydroxybutyrate (BHB), the most prevalent circulating ketone body across all groups in both males and females. During fasting, we observed a significant increase in BHB in mice fed LFD, LFMP and HFD in both males and females (**Figure 1C,1D**). However, in KD fed mice, the levels of BHB were elevated regardless of feeding:fasting status and, furthermore, significantly higher than any other group in the fasted state. As another way of evaluating the ketogenic nature of our diet, we placed males on LFD, HFD and KD in an indirect calorimetry system to interrogate their substrate usage through their respiratory exchange ratio (RER). RER records shows that mice on LFD alternate between more carbohydrate burning during the feeding phase to more fat burning during the fasting state (**Figure S1A**). In contrast, HFD RER oscillations are severely dampened with an average value of 0.78. Mice on KD display a flat RER with even lower RER values of 0.73 suggestive of constant ketosis regardless of time-of-day or feeding status (**Figure S1A**). Together, these data suggest that the KD used in this study produces a constant state of ketosis and elevated circulating levels of BHB.

**Figure 1.**
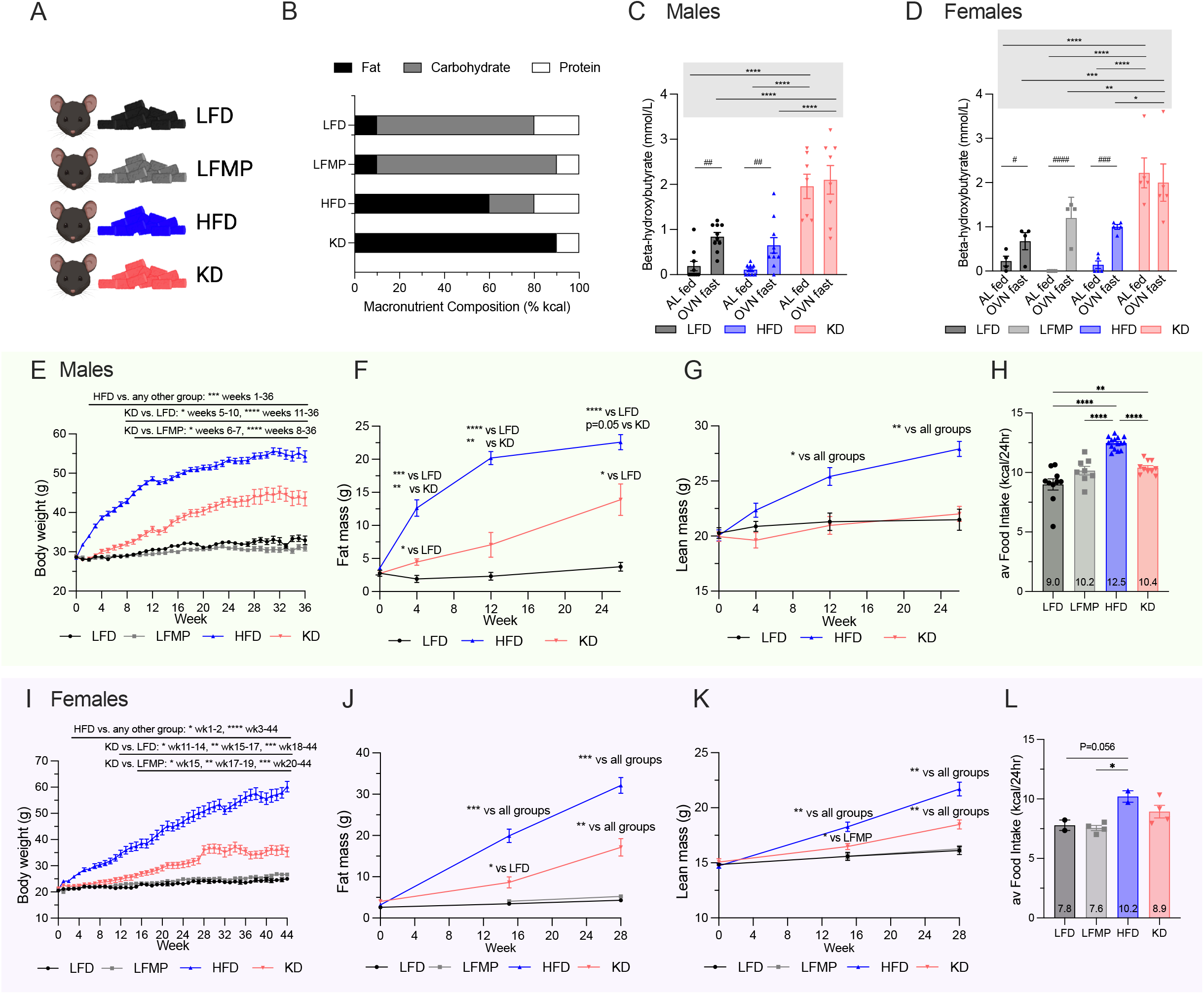
Mice on KD are protected from weight gain as compared to mice on HFD. A) Diets and color scheme used throughout the paper. B) Macronutrient composition of diets used. C) Male fed and fasted beta-hydroxybutyrate levels taken at 4 weeks (overnight fast) and 6 weeks (*ad libitum* fed); n=7-10/ group. D) Female fed and fasted beta-hydroxybutyrate at 15 weeks on diets (n=4-5/ group). E) Male body weight pooled from 3 independent cohorts (n= 30-50/ group). F, G) Male body composition (n=5-6/ group): F) fat mass, G) lean mass. H) Average 24-hour food intake from 36 weeks of data collection from 3 independent cohorts (n= 8-14cages /group). I) Female body weight pooled from two independent cohorts (n=10-20/ group). J, K) Female body composition (n=5-10/ group): J) fat mass, K) lean mass. L) Average 24-hour food intake from 26 weeks of data collection from 2 independent cohorts over (n=2-4 cages/ group). <u>Statistics</u>: One way ANOVA was used for H and L, and mixed effects ANOVA was used for all other comparisons, post Hoc testing used Tukey’s HSD test. * p<0.05, ** p<0.01, ***p<0.001, **** Pp<0.0001, # only used for within group comparisons, #p<0.05, ## p<0.01, ### p<0.0001. Data are represented as mean ± SEM.

Although the KD is 89.9% fat, male and female C57Bl/6J mice on KD were prevented from weight gain as compared to mice on a 60% HFD (**Figure 1E,1I**). Both males and females on 60% HFD rapidly gained weight. BW divergence between the HFD and the low-fat diets started as early as 1 week in both sexes. BW gain on KD was slower and diverging from low-fat diets only after over 10 weeks in males (**Figure 1E**) and 11 weeks in females (**Figure 1I**) under the different diet regimen. BW remained significantly lower in KD than HFD at all times (**Figure 1E,1I**). At the end of 36 weeks, male mice on HFD weighed 54g versus 43g on KD (**Figure 1E**). In female mice, after 44 weeks, mice on HFD weighed 60g versus 35g for mice on KD (**Figure 1I**).

In males, weight gain on KD was mainly due to changes in fat mass as no changes in lean mass were observed across the 26 weeks (**Figure.1F,G**). This is different to mice on HFD, in which weight gain was also predominantly due to increased fat mass, but changes in lean mass were also observed with mice on HFD gaining 19g of fat mass and 8g of lean mass after 26 weeks (**Figure 1F**). Further, in HFD-fed males, fat mass accrual was maximal after 12 weeks, and plateaued between weeks 12 and 26, but lean mass increased throughout the whole study (**Figure 1F,1G**). In both males on HFD and KD, fat mass was higher than in the LFD group from week 4 at any time onwards. However, lean mass was only greater in the HFD, and lean mass not different between LFD and KD. After 26 weeks on the diet, although the mice on HFD had more fat and lean mass, the percentage of body fat was not different between HFD- and KD-fed males, 42.7 ±1.6% fat for HFD 35.7 ±5.1% fat for KD (**Figure S1B**).

In females, weight gain on a KD was due to increases in both lean and fat mass with mice gaining roughly 3.5g lean mass and 13g fat mass (**Figure 1J,1K**). Similar to the males, females on HFD gained most of their weight through increases in fat mass but had a higher proportion of weight gain coming from fat with a roughly 29.5g increase in fat mass and 7g increase in lean mass (**Figure 1J,1K**). After 27 weeks, mice on HFD had 63.7 ±1.2% body fat versus 46.7 ±3.7% fat on KD (**Figure S1**).

The differences in weight between groups can at least partially be explained by differences in food intake (**Figure.1H,1L**). Average food intake in male and female mice was highest in mice on HFD followed by mice on KD. This difference between HFD and KD was significant in male mice with mice on HFD consuming 13.4 kilocalories (kcals) versus 11.1 kcals on KD and 8.4 kcals and 10.3 kcals for LFD and LFMP respectively (**Figure 1H**). Furthermore, throughout 36 weeks of intake measurements, the mice on HFD consumed more than the other groups, although this difference was more pronounced early on, it remained for the duration of the study (**Figure S1D**). Throughout the study, mice on KD also consumed an intermediate amount of calories that was higher than the low-fat diet groups and lower than the HFD group (**Figure S1D**). In contrast to the males, female mice consumed less food overall with mice on HFD consuming 10.2 kcals versus 8.9 kcals on KD and 7.8 kcals and 7.6 kcals for LFD and LFMP respectively. However, in female mice the only significant difference in food intake was between the HFD and LFMP with HFD versus LFD having a marginally significant difference. This lack of significance between HFD and KD and HFD and LFD is likely because of the low cage number since food intake was measured as average cage intake each week (**Figure 1L**). There were no differences in food intake between the low-fat diet groups neither in males nor in females (**Figure 1H,1L**). During the first week that the females were on the intervention diets, the HFD group consumed significantly more than the other groups, and while this difference declined, the HFD group still consumed more than other groups throughout the study (**Figure S1D**).

### 2 KD-fed mice have severe plasma hyperlipidemia and males have liver steatosis and dysfunction

Because KD contains ∼90% of fat, we assessed lipid homeostasis in KD-fed mice. In both males and females, plasma triglycerides (TG) were significantly higher in KD-fed mice than in any other group including HFD-fed mice (**Figure 2A,2B**). In general, TG levels were lower in females than males, yet the TG increase in response to KD was similar in both sexes wherein TG levels were 1.7 times higher in KD than HFD fed mice (336 vs 193 in males and 152 vs 92 in females). Similar to the changes in TG, plasma non-esterified fatty acid (NEFA) levels were significantly higher in KD-fed mice versus other diet groups– 2.75 and 1.8 times higher in KD vs. HFD in males and females respectively (**Figure 2C,2D**). There were not sex-differences in NEFA levels (**Figure.2C,2D**). Plasma cholesterol (Cho) responses to the diets were different than that of TG and NEFA, with similar elevations in Cho levels in both HFD and KD-fed mice as compared to mice on either of the low-fat diets (**Figure S2A,S2B**) [sex difference in Cho levels: Cholesterol levels were higher in all groups in male mice as compared to female mice]. Together we found, that in contrast to HFD, KD significantly raises plasma TG and NEFA levels in both males and females mice suggesting an overabundance of fat is still present in the body even though mice on KD are leaner than mice on HFD.

**Figure 2.**
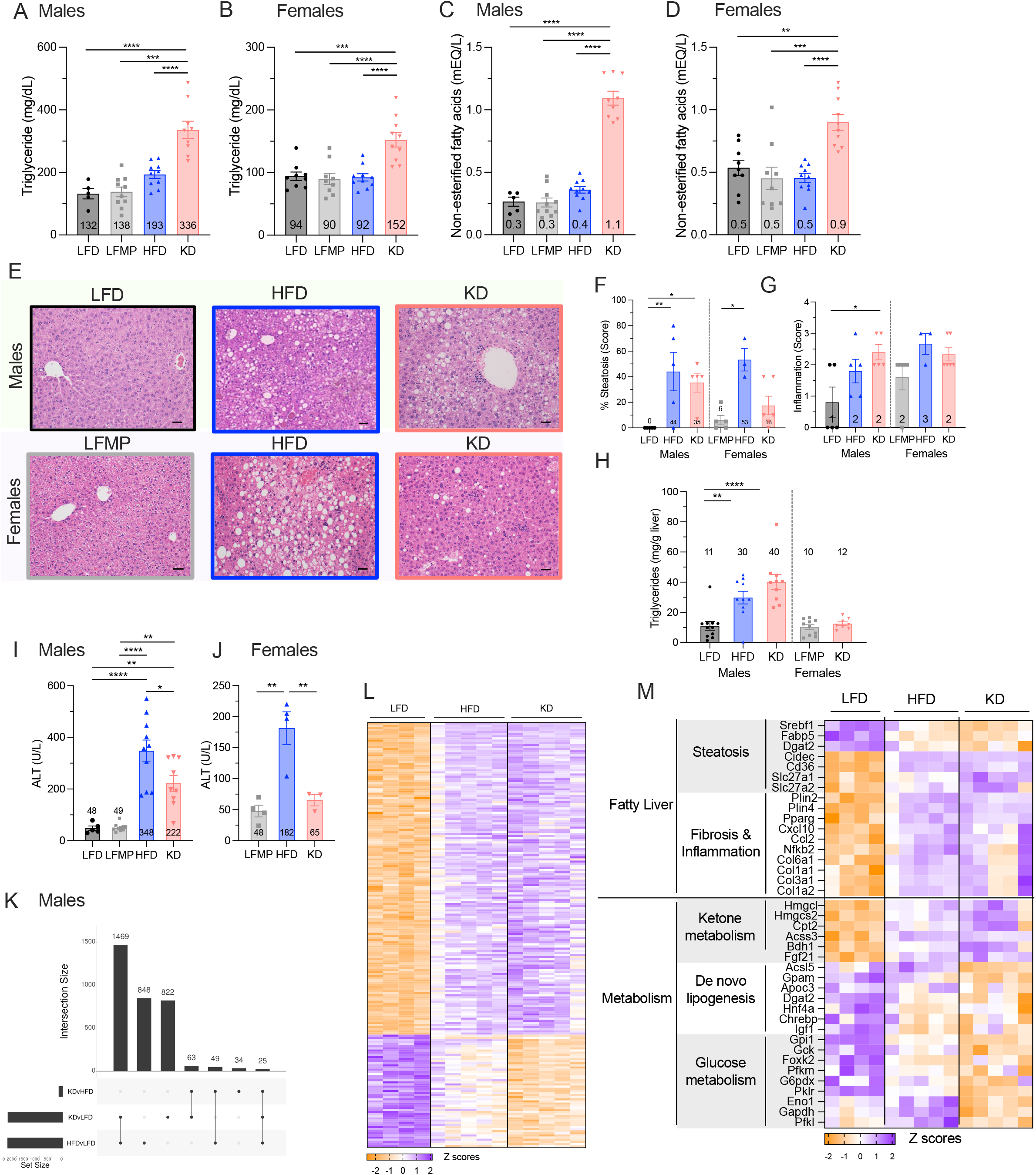
A long-term KD causes dyslipidemia regardless of sex, and males have steatosis and liver dysfunction. A, B) Triglyceride and (C,D) non-esterified fatty acids (NEFA) plasma levels in (A,C) male mice after 32 weeks on diets (n=5-10/ group), and (B,D) female mice after 15 weeks on diets (n=9-10/ group). All blood samples were collected in the fed state in the middle of the dark phase. E) Representative images from male mice on diets for 28 weeks (top row) or female mice on diets for 33 weeks (bottom row). Scale bars are 20μm. F, G) Results of liver scoring by a liver pathologist blinded to conditions (male n=5/group; female n=3-6/ group, HFD females were part of a different cohort). F) Steatosis as estimated percent visual field containing fat. G) Inflammation score from 0-3 based on average number of inflammatory foci per visual field with five visual fields scored as described by Liang et al^85^. H) Liver triglyceride content in males from two independent cohorts on diets for 28 weeks and 38 weeks (n= 10-11/ group) and females (n= 8-10/ group) on diets for 33 weeks. I, J) Plasma alanine aminotransferase (ALT) levels in males (I) after 32 weeks on diets (n=5-10/ group) and females (J) after15 weeks on diets (n=9-10/ group). K-M) Liver bulk RNA sequencing results. Livers were harvested from male mice after 38 weeks of their respective diets n=4-5/ group. K) UpSet plot comparing all differentially expressed genes. L) Heatmap of the top 200 most significant differentially expressed genes between LFD and KD. M) Heatmap of selected genes associated with steatosis, fibrosis, and metabolism. (L, M) Color intensity in the heatmaps represents the Z score based on Log2 expression values. <u>Statistics</u>: A-F, H-J) Data are represented as mean ± SEM. One-way ANOVA with Tukey’s post hoc testing with p values as: * p<0.05, ** p<0.01, ***p<0.001, **** p<0.0001. G) Kruskal-Wallis test with Dunn’s multiple comparison tests for post hoc analyses. K-M) RNAseq statistics were determined using DESeq2 analysis with a Wald test and Benjamini-Hochberg correction. Significant genes had an adjusted p-value of <0.05.

The hyperlipidemic profile of KD-fed mice prompted us to assess liver health and function in male and female mice. Hematoxylin and eosin-stained (H&E) sections of livers collected after six months of dietary interventions, were examined by a board-certified liver pathologist blinded to the feeding condition. In males, steatosis, as estimated by the percentage of area made up of lipid droplets, was higher in mice on HFD and KD as compared to mice on LFD and was not different between HFD and KD-fed mice (**Figure 2F**). In females, only the livers from HFD mice had significant steatosis with no differences between LFMP and KD-fed females (**Figure 2F**). The visual inflammation scoring did not reveal dramatic or significant differences between any groups other than a possible trend to higher inflammation in both HFD and KD-fed mice in both sexes (**Figure 2G**). These results suggest that from a histology standpoint, the livers from HFD and KD male mice were very similar, with ectopic lipid deposition occurring in both, whereas TG did not accumulate in female livers on KD. These results were confirmed using biochemical quantification of hepatic TG content (**Figure 2H**). Male mice on long term 6-9 months KD and HFD had significantly elevated liver TG content as compared to mice on LFD (3 to 4 times higher), whereas female mice on KD did not have elevated liver triglycerides (**Figure 2H**). Together these results suggest that steatosis is similar between males and females on long-term HFD but only appears in males under KD, suggesting sex differences in the liver response to KD. Finally, we measured plasma alanine transaminase (ALT) as a readout of liver function. In males, after 32 weeks of their respective diets, HFD- and KD-fed mice had roughly 7- and 4.5-times higher ALT compared to low-fat diets respectively (**Figure 2I**). In addition, ALT was significantly lower in KD vs. HFD males. In females, after 48 weeks on their respective diets, mice on HFD and KD also had elevated ALT. ALT was not different between mice on LFD and KD, but the HFD group had ALT levels that were 3.8 and 2.8 times the levels of mice on LFMP or KD respectively (**Figure 2J**). Overall, these findings indicate liver dysfunction and steatosis under KD feeding in males but not females suggesting that there are sex differences in the liver response to KD.

To evaluate how HFD versus KD impacted hepatic lipid handling at the molecular level, we compared the liver transcriptome of KD-, HFD- and LFD-fed males. Gene expression profiles were very similar between mice on HFD and KD, with only 171 differentially expressed (DE) genes (adj p<0.05) between mice on KD and HFD compared to 2351 for HFD versus LFD and 2371 for KD versus LFD (**Figure 2K**). The gene expression profiles of the top 200 DE genes between KD and LFD were virtually identical for HFD and KD (**Figure 2L**) and many of these genes are associated with steatosis, fibrosis and inflammation in agreement with our histology and biochemical findings. (**Figure 2M**). Thus, at the gene expression level, while both HFD and KD livers are different from mice on LFD, the livers from mice on HFD and KD are almost indistinguishable. Because of the constant state of ketosis observed in KD mice, we specifically interrogated the expression of genes involved in ketogenesis and glucose metabolism. Genes linked to ketone bodies metabolism – such as 3-hydroxy-3-methylglutaryl-CoA synthase 2 (*Hmgcs2),* controlling the rate limiting state in ketogenesis, and β-hydroxybutyrate dehydrogenase 1 (Bdh1)-were as expected increased under KD, yet, interestingly similar expression patterns were also observed in HFD (**Figure 2M**). Additionally, genes involved in de novo lipogenesis such as *Chrebp* and *Dgat2* were downregulated in both HFD and KD livers as compared to LFD (**Figure 2M**). Expectedly, genes related to glycolysis and glucose metabolism were lower in KD as compared to either HFD or LFD (**Figure 2M**). Overall, liver histological, biochemical and molecular analysis revealed that, although mice on a KD are leaner than mice on a HFD, the males still suffer steatosis and inflammation that likely impair their liver function.

### 3 Mice on KD have severe glucose intolerance and impaired insulin secretion

We next examined how a very high fat low carbohydrate KD affects glucose homeostasis compared to a traditional 60% HFD and low-fat diets. In the first four weeks of the intervention, mice on KD had lower fasting blood glucose levels compared to HFD, as well as lower postprandial glucose compared to both HFD and LFD (**Figure 3A**). However, when tested after a longer time, between 27-32 weeks for males and 15 weeks for females, fasting glucose was similarly elevated in HFD- and KD-fed mice compared to low fat diets (**Figures 3B,3C**). Fed blood glucose remained significantly lower in KD-fed than in HFD-fed males and females and than LFD in males likely due to the very low amount of carbohydrates in the diet (**Figure 3B**). These results suggest that while short term, a KD may be beneficial for regulating (hyper)glycemia, long term KD feeding is associated with fasting hyperglycemia, similar to that observed under HFD and fed hypoglycemia. We next assessed the ability of the mice to restore glycemia after an intraperitoneal (IP) bolus of glucose. IP Glucose Tolerance Test (GTT) revealed glucose intolerance similar to HFD-fed mice after 11 weeks in males and 31 weeks in females (**Figure 3D,3F**). However, much more severe glucose intolerance was observed in males after 33 weeks on the diet: KD males reached a maximum of 539 mg/dL of glucose versus 427 in HFD, and blood glucose only decreased by 95 mg/dL versus 168 mg/dL in HFD after 2 hours; despite KD mice receiving less glucose based on their lower body weight (**Figure 3E**). Oral glucose tolerance tests as well as oral mixed meal tolerance tests also confirmed severe glucose intolerance in mice on KD (**Figure S3A-C**). Taken together, these findings show that male and female mice on long-term KD develop glucose intolerance.

**Figure 3.**
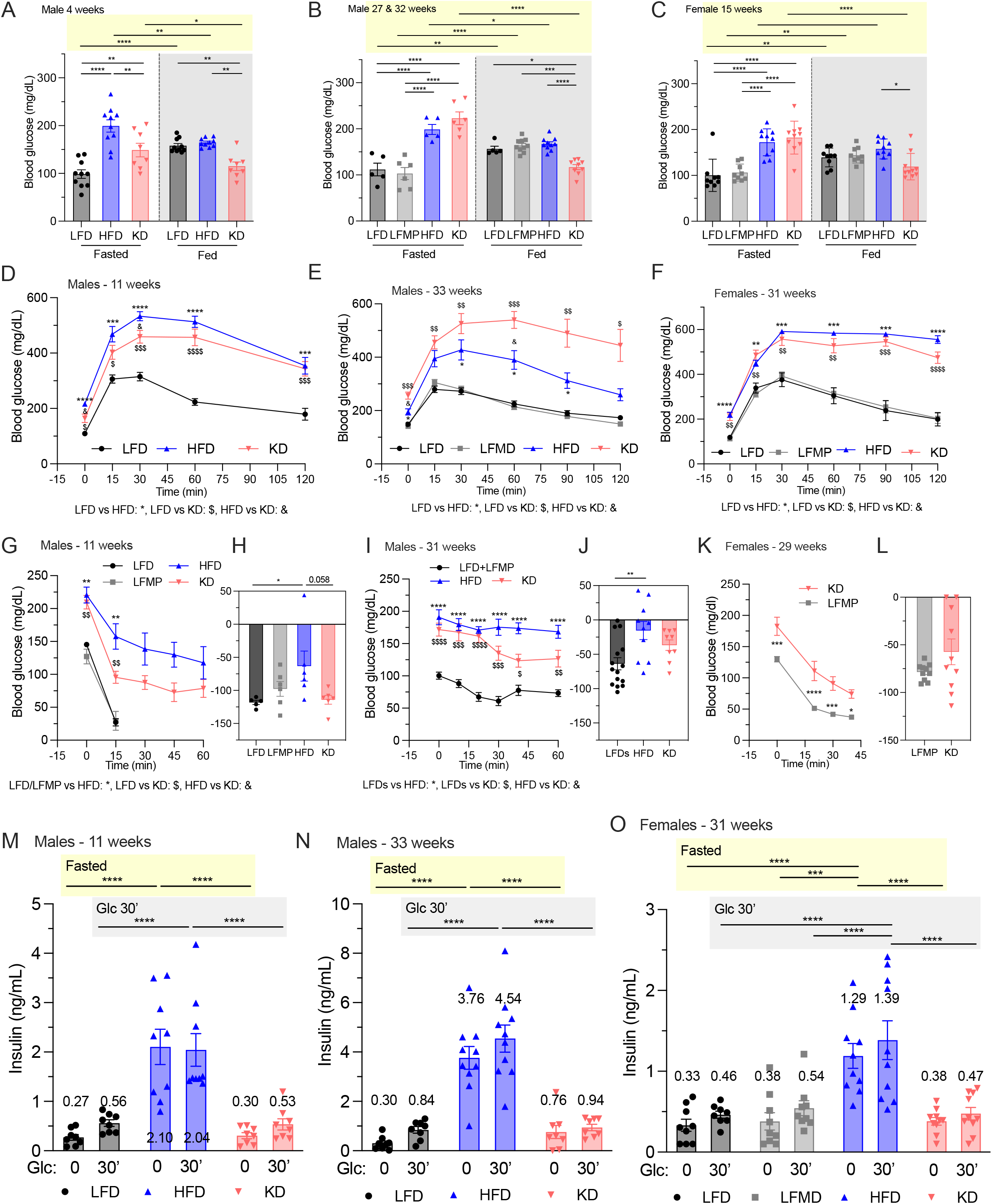
A long-term KD causes severe glucose intolerance and suppresses insulin secretion. A-C) Fasted and fed blood glucose. Fasted glucose was measured at 7am (onset of dark cycle) following a 14 hour fast, and fed glucose was measured at 11am during the dark cycles. A) Males 4 weeks (n=8-10/ group), B) Males 27 and 32 weeks from two independent cohorts (n=5-10/ group). C) Females after 15 weeks on diets (n=4-9/ group). D-F) Glucose tolerance tests with a bolus of 1.5mg/g glucose given after an overnight fast (over the light phase) in: D) Males after 11 weeks on diets (n=8-10/ group), E) Males after 33 weeks on diets (n=5-6/ group), and F) females after 31 weeks on diets (n=8-10/ group). G-L) Insulin tolerance tests with 0.75IU/kg insulin. Bar graphs (H, J, L) show the slopes in the first 15 (H) or 30 minutes (J,L). G-H) Males after 11 weeks on diets (n=5/ group). I-J) Males on diets for 33 weeks (n=5-10/ group). K-L) Females on diets for 29 weeks (n=5-10/ group). M-O) Insulin levels taken at baseline and 30 minutes after a bolus of glucose. Taken during the same experiments as D-F). <u>Statistics</u>: One-way or two-way ANOVA with Tukey’s HSD for post hoc testing was used. The number of symbols represent p values as: * p<0.05, ** p<0.01, ***p<0.001, **** p<0.0001. Data are represented as mean ± SEM.

To test whether glucose intolerance in KD-fed mice was due to insulin resistance, we performed insulin tolerance tests (ITT). After 11 weeks on the diet interventions in males, fasted blood glucose (BG) was similar between HFD and KD groups and significantly higher than LFD groups (**Figure 3G**). Upon insulin injection, BG dropped drastically within the 15 minutes with a delta or slope that was not different between KD and the LFDs groups (-118.2mg/dl in LFD, - 97.7mg/dl in LFMP, -114.0mg/dl in KD) but significantly less in HFD (-63.2mg/dl in HFD)(**Figure 3G,3H**). A similar response was observed in females at 29 weeks (**Figure 3K,3L**). When an ITT was performed in males after at 32 weeks on their respective diets, the response to insulin was maintained in KD and LFD mice while the HFD-fed males hardly responded (**Figure 3I,IJ**). Together these results confirmed the well-characterized insulin resistance of males fed HFD but suggested that KD-fed male and female mice were not insulin resistant.

To determine whether glucose intolerance in KD-fed mice was instead due to a defect in insulin secretion, we measured plasma insulin levels in response to a glucose bolus using an *in vivo* glucose stimulated insulin secretion (GSIS) assay. We found that in males and females on LFDs and KD, baseline fasted insulin levels were low and not different, in contrast to the HFD groups which had high insulin both prior and 30’ after the glucose injection, as would be expected with insulin resistance (**Figure 3M-3O**). LFD and LFMP fed mice had a modest yet significant increase in insulin upon glucose injection which was sufficient to restore normoglycemia in about 1h (see GTTs **Figure 3D-3F**). The glucose bolus did not lead to a rise in insulin levels in KD-fed mice, corresponding to the lack of BG regulation seen in the GTTs.

Together these results suggest that altered glucose homeostatic response in HFD-versus KD-fed mice are not caused by the same mechanism. While impaired glucose regulation in HFD-fed mice is likely linked to insulin resistance in the context of high insulin levels, impaired insulin secretion likely underlies glucose intolerance in KD-fed mice. A time-course experiment further revealed that insulin secretion deficit occurred after 4-8 weeks of KD (**Figure S3G-I**) suggesting that KD impairs islets function over long-term feeding.

### 4 KD-fed mice have normal islets size and insulin content

Since insulin secretion was impaired in mice on KD, we next used histologic analysis to determine whether pancreatic islet size or number were reduced in mice on a KD. Since insulin secreting β cells make up the majority of cells in the islets in mice, we used insulin/ proinsulin-stained area to demarcate islets by immunochemistry. The average islet area was much larger in HFD-than KD- or LFD-fed mice but there was no difference in size in KD compared to LFD islets (**Figure 4A,4D**) while the number of islets per section was similar in all three groups (**Figure 4C**). When examining islets size distribution by group, we also found that mice on HFD had significantly fewer small islets and at least twice more larger islets compared to the other groups (**Figure 4B**, **Figure S4B**). We also found that the percent insulin positive area was not different between LFD and KD but was higher in the HFD group (**Figure S4A**). Mirroring the findings in males, there were no differences in average size, size distribution, or number of islets between females fed a KD or a LFMP (**Figure 4E,4F,4G**). Islet size and number were also comparable between males and females. To complement these histological findings, we quantified pancreas mass and insulin content. Pancreas mass was significantly increased in both HFD-fed males and females (**Figure 4H,4J**) but was similar between mice on LFD and KD (**Figure 4H,4J**). In agreement with our histological findings, HFD pancreas had more insulin per milligram tissue in both male and female mice (**Figure 4I,4K**), while male mice in general had more insulin than female mice. The differences in pancreas size and insulin content also translated to the HFD having more total insulin per pancreas as compared to the KD and LFD mice (**Figure S4C,S4D**). Overall, these results confirm the known effect of HFD-feeding on islets hyperplasia and show that KD feeding was not associated with overt loss in islet mass or insulin content that could underly insulin secretion deficiency.

**Figure 4.**
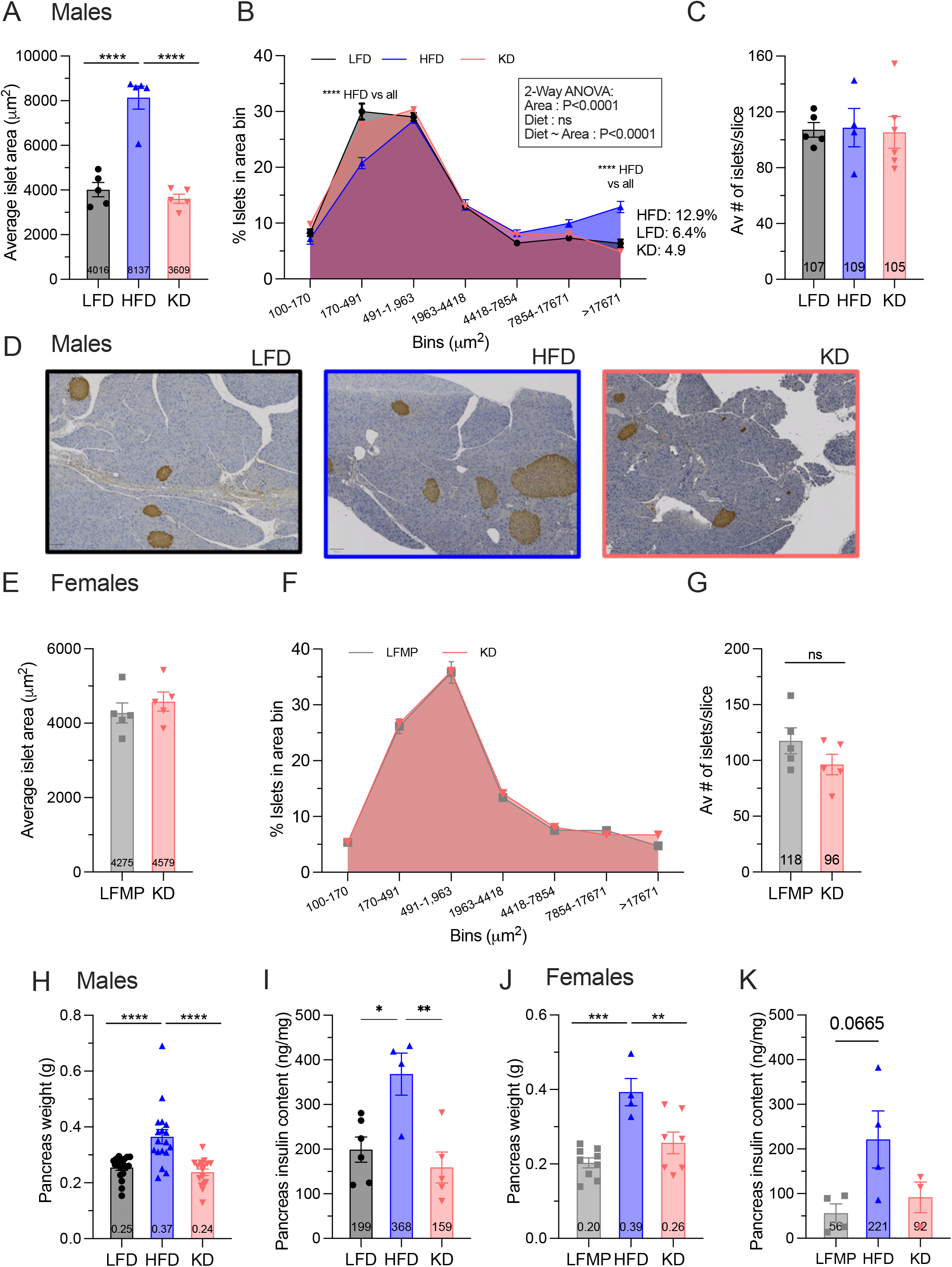
A HFD but not a KD increases islet size, pancreas weight, and insulin content in the pancreas. A-G) Analysis of chromogenic insulin/proinsulin staining in the pancreas with hematoxylin counterstain. Qupath was used to quantify islet area based on detection of the brown insulin stain. A-C) Males, 8-12 slices were taken throughout the pancreas from n=4-6 animals per groups for a total 5157 islets analyzed on LFD, 6718 on HFD, and 6389 on KD. E-G) Females, 6-7 slices were taken throughout the pancreas (n=5/ group), for a total of 3749 LFMP and 3428 KD islets analyzed. A, E) Average islet area. B, F) Frequency distribution of islet sizes as number of islets per slice in each bin. Bins correspond to islet diameter 100μm^2^= diameter of 11.3μm or the size one beta cell (and the minimum size needed to identify a nucleus and clear borders in the analysis), 170μ^2^= diameter of 14.7μm, 490.87μm^2^= diameter of 25μm, 1963μm^2^= diameter of 50μm, 4417.86μ^2^= diameter of 75μm, 7853.981μm^2^= diameter of 100μm, 17671μm^2^= diameter of 150μm. C, G). Average number of islets per slice. D) Representative images from male islets. Scale bar is 100μm. H) Male pancreas weight after 6-9 months on respective diets (n=17-18/ group pooled from 3 independent cohorts) and (I) insulin content after 6 months on the diets (n=5-6/ group). J) Female pancreas weight (n=4-8/ group) and (K) insulin content after 6 months on respective diets n=3-4/ group. <u>Statistics:</u> One-way or two-way ANOVA with Tukey’s HSD for post hoc testing was used. The number of symbols represent p values as: * p<0.05, ** p<0.01, ***p<0.001, **** p<0.0001. Data are represented as mean ± SEM.

### 5 Mice on KD lose rapid insulin secretion *in vivo* and isolated islets have impaired insulin secretion

To delve further into the underlying cause of glucose intolerance in KD-fed mice, we performed an *in vivo* hyperglycemic clamp experiment in awake, freely moving mice. Throughout the 120-minute protocol, 50% glucose was infused through a jugular vein catheter to obtain 30 minutes of clamped BG at around 250mg/dL for 30 minutes. To that end, BG was measured at regular intervals throughout, and the infusion rate titrated accordingly. Upon initial infusion, BG rapidly raised in all groups and peaked at 30 min (**Figure 5A**). The Glucose Infusion Rate (GIR) was progressively reduced as BG declined to the target 250 mg/dL, and the GIR was then progressively increased after about 60 min to maintain the clamped value which was obtained in LFD and HFD-fed mice between minutes 90-120 of the test (**Figure 5C**). In KD-fed males however, BG remained significantly elevated for 90 minutes despite the GIR being reduced to the minimum infusion rate possible from 50 minutes onwards (**Figure 5C**). BG in KD mice only started to decrease toward the end of the experiment (**Figure 5A**). From 90-120 minutes, BG was effectively clamped at an average of ∼230 mg/dL in HFD and LFD fed mice, but BG in KD-fed mice was significantly higher at 314 mg/dL (**Figure 5B**) even in the context of significantly lower GIR (**Figure 5D**). In parallel, we measured plasma insulin levels throughout the study (**Figure 5E**). Upon glucose infusion, insulin levels significantly and rapidly rose in LFD and HFD (87% and 68% increase resp.), reaching max level at 15 min (**Figure 5E,5F**). Strikingly, insulin levels in KD-fed mice initially decreased from 0-10 minutes; a slow rise in insulin levels wasn’t apparent until 30 min post GI start; and insulin levels peaked at 90 min which coincide with the time at which blood glucose begins to decline (**Figure 5A**). Insulin levels overall were higher in HFD than LFD during the clamp (**Figure 5E**) which is indicative of insulin resistance. Since mice on KD have similar insulin levels to the control mice both at baseline and at the peak, this suggests that their glucose intolerance is caused by impaired insulin secretion rather than insulin resistance.

**Figure 5.**
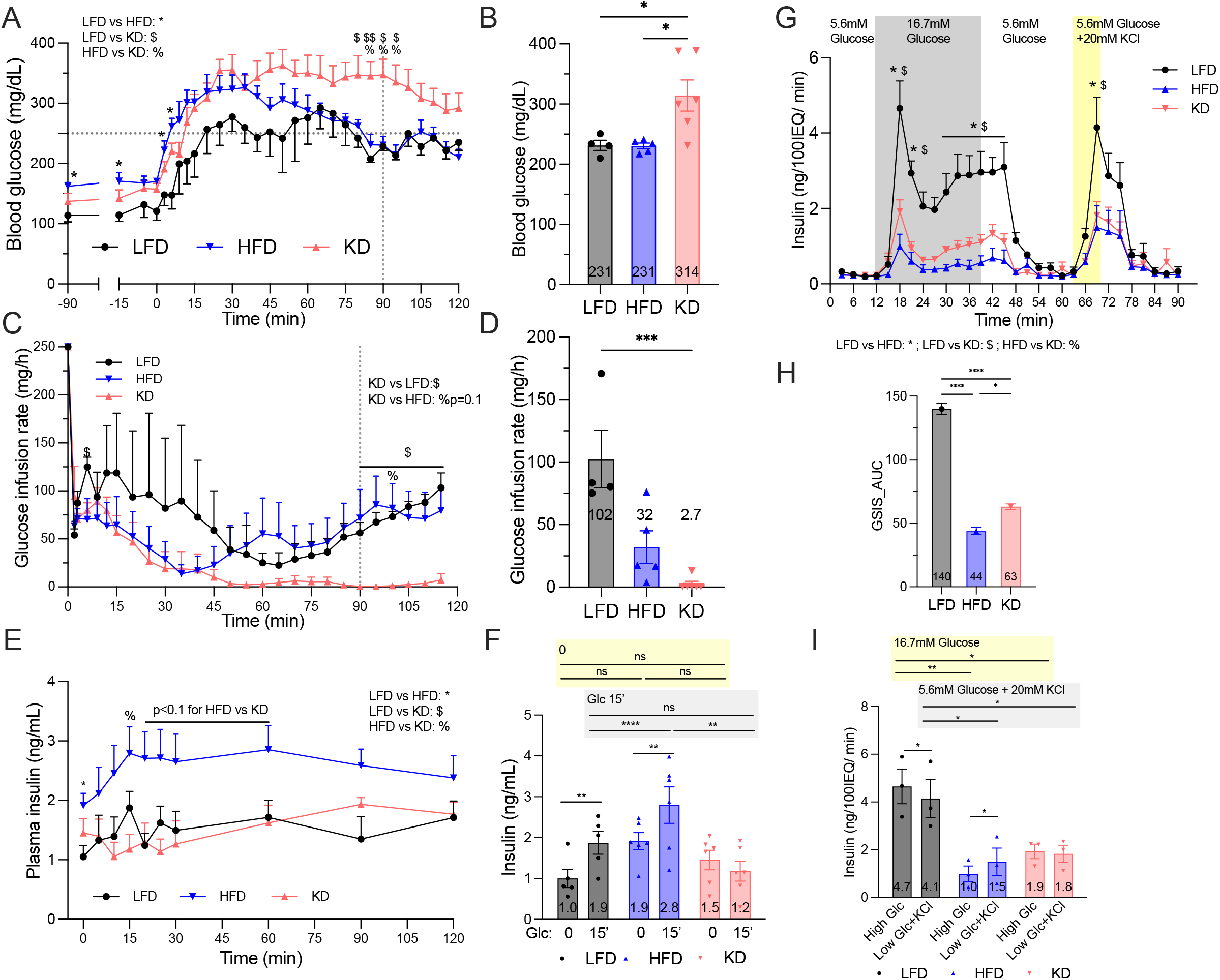
Mice on KD lose rapid insulin secretion. A-F) Measures taken during hyperglycemic clamp after 25 weeks on the respective diets. n=4-6 male mice per group. Time courses of blood glucose (A), the glucose infusion rate (C), and plasma insulin levels (E) obtained via tail vein sampling. Average blood glucose (B) and average glucose infusion rate (D) during the 90-120 minutes when blood glucose was clamped. F) Insulin levels before and 15min into the glucose infusion. G-I) Ex vivo GSIS results (n=3/group). Insulin secretion is normalized to 100 insulin equivalents (IEQ; 1 IEQ represents an islet with a diameter of 150μm). H) Area under the curve for insulin secretion during the perifusion. I) Peak insulin secretion during high glucose and KCl stimulation. <u>Statistics</u>: One- or two-way ANOVA with post hoc testing using Tukey’s HSD except in H which used Dunnett’s multiple comparison test to compare HFD and KD with the LFD group only. The number of symbols above data denote the following p values: * p<0.05, ** p<0.01, ***p<0.001, **** p<0.0001. Data are represented as mean ± SEM.

To assess cell-autonomous differences in insulin secretion, islets from males fed LFD, HFD and KD for 60 weeks were isolated and tested by dynamic *ex vivo* GSIS using a perifusion system (Vanderbilt Islet and Pancreas Analysis core). Baseline insulin secretion at 5.6mM glucose (roughly equal to 100mg/dL) was similar across all three groups. However, upon stimulation with 16.7mM glucose (roughly 300mg/dL), both KD and HFD islets had less insulin secretion than LFD islets (**Figure.5G, 5G**). Specifically, the initial peak in insulin secretion observed six minutes after the introduction of high glucose (minute 18) in LFD was blunted in both HFD and KD islets (4.7 compared to 0.99 for HFD and 1.9 ng/100IEQ/min for KD), and insulin levels remained lower throughout the high-glucose stimulation. In the final step of the GSIS, islets were exposed to 20mM potassium chloride (KCl) to depolarize b-cells membranes and assess maximal insulin secretory capacity independent of glucose metabolism. Both HFD and KD islets responses were dampened compared to LFD islets (**Figure 5G**). Peak insulin secretion in HFD-islets under KCl was higher than under high glucose (1.5 vs 0.99 ng/100IEQ/min), suggestive of a deficiency in both the glucose sensing and the insulin secretion machinery. Under KD, the peak response under KCl and high glucose were similar (1.82 and 1.9 ng/100IEQ/min), suggesting a deficiency in insulin secretion machinery. Taken together, the results of dynamic GSIS testing suggest that insulin secretion is compromised/impaired in both HFD and KD compared to LFD, possibly through different mechanisms.

### 6 Transcriptomic and EM analysis of KD islets reveals ER-Golgi stress and aberrant Golgi function

To gain insight into the mechanisms behind the secretory dysfunction of KD islets and how they may be different to HFD islets, we analyzed the bulk transcriptome of isolated islets harvested from male mice fed LFD, HFD, or KD for 36 weeks. We found the most gene expression changes when comparing islets from mice on KD to those from mice on LFD with a total of 1666 differentially expressed (DE) genes. Amongst those, 1083 were uniquely changed between KD and LFD, 397 distinguish KD versus both HFD and LFD, 175 changed in both KD and HFD compared to LFD, and a very small 11 genes were different between all dietary condition (**Figure 6A**). In contrast, there were only 242 differentially expressed genes in HFD vs LFD (**Figure 6A**), indicating that the KD altered the islet transcriptional profile much more than a HFD when compared to LFD. Finally, comparing KD to HFD, there was a total of 1127 DE genes. Amongst those, 697 were changed only under KD vs HFD whereas 397 changed in both KD compared to HFD and LFD, and a small 22 DE genes changed between KD vs HFD & HFD vs LFD)(**Figure 6A**). Overall, transcriptomics analysis thus revealed ∼400 DE genes that distinguish the islets from KD-fed mice from islets of mice fed either HFD or LFD (397 + 11) (**Figure 6A**).

**Figure 6.**
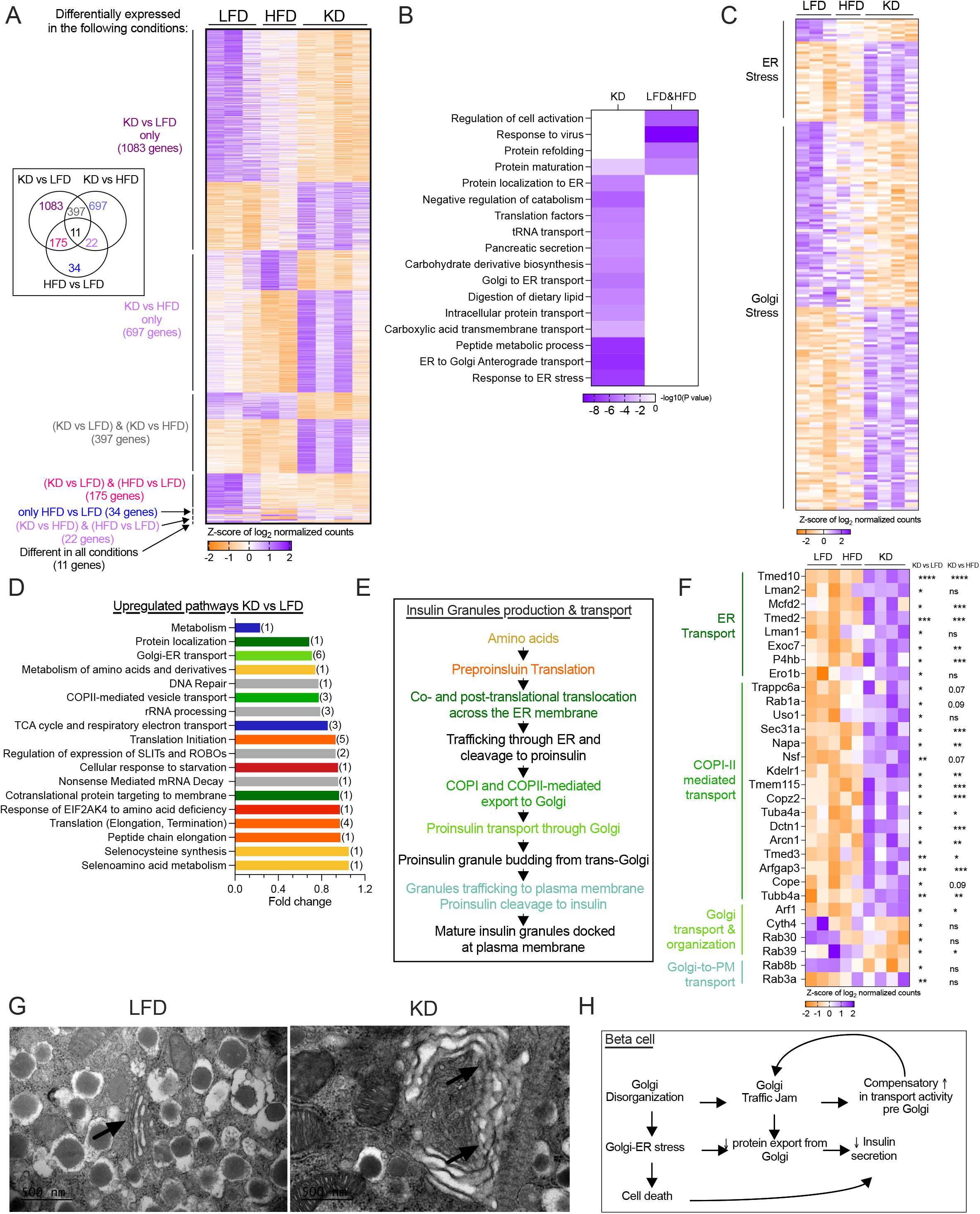
Islet transcriptomic analysis reveals that KD causes ER Golgi stress and impaired protein transport. Bulk RNA sequencing from islets isolated from male mice after 36 weeks on interventions. A, C, F) Heatmaps showing Z scores of log2 normalized expression. A) Heatmap of all differentially expressed genes. The Venn diagram inset shows number of genes per comparison, and the colors correspond to the colored labels on the heatmap. B) the 397 genes that were differentially expressed in both KD vs LFD *and* KD vs HFD were run through Metascape which identified ontology categories that were enriched in one group versus the other as displayed in the heatmap; the purple indicates increased expression in the corresponding group, and the color intensity represents -log10(P-value) of the enrichment as calculated by Metascape. C) Using the Gene Ontology database, we identified genes involved in ER and Golgi stress genes, and plotted the genes that were significantly different in KD versus LFD. Heatmap of selected genes associated with ER-Golgi stress D) Significantly upregulated pathways identified using Reactome Pathway analysis of differentially expressed genes between KD and LFD. FDR <0.05 for all pathways. E) Pathway of insulin granule generation and processing. Colors in E correspond to colors in D and F. F) Heatmap of selected genes involved in protein processing and vesicular transport. G) Electron micrograph images of pancreatic beta cells showing Golgi dilation in KD. H) Mechanism linking transcriptomic findings with reduced insulin secretion. <u>Statistics</u>: Differentially expressed genes were identified using DESeq2 analysis with a Wald test and Benjamini-Hochberg correction. Significant genes had an adjusted p-value of <0.05.

To identify potential mechanisms behind KD-specific islet secretory dysfunction, we performed pathway enrichment analysis using this list of DE genes and Metascape. Pathways linked to translation and peptide metabolism, ER stress, and ER to Golgi transport were specifically enriched in KD versus LFD and HFD (**Figure 6B**). Additionally, the most significantly upregulated gene in KD as compared to both HFD and LFD was *Creb3l3* (Figure S5A,S5B), a member of the CREB3 transcription factor family that is activated by ER and Golgi stress.^32^ Since these analyses pointed at a potential ER and Golgi stress signature, we therefore performed a targeted analysis of genes related to “Golgi Apparatus” (GO:0005794) and “Response to Endoplasmic Reticulum Stress” (GO:0034976) (**Figure 6C**). 158 out of 1280 genes (12.3%) linked to Golgi and almost 20% (41 out of 224) of GO annotated ER stress genes showed significantly altered expression in KD islets versus LFD or HFD islets (**Figure 6C**). This targeted analysis further evidenced severe ER-Golgi stress in the KD islets. Zooming out, we used the bigger list of KD vs LFD 1666 DE genes and Reactome to pathway analysis. Reactome identified significantly upregulated pathways in KD linked to (i) translation (orange, 10 pathways), (ii) metabolism (blue, 4 pathways), (iii) Golgi-ER transport and COPII-mediated vesicle transport (green, pathways), (iv) protein localization (dark green, 2 pathways), and (v) amino acid metabolism (yellow, 3 pathways). Similar enrichment of genes related to vesicle transport, ER/Golgi stress, and translation was also observed when KD and HFD islets were compared (**Figure S5C**). A color-coded schematic (**Figure 6E**) shows that these altered pathways can influence insulin secretory capacity throughout the process of insulin synthesis and granules secretion. Targeted investigation/Further probing the expression of genes involved into genes that are specifically involved in vesicle transport between the ER and Golgi and to the plasma membrane (as identified by Reactome and literature search) shows that many of these genes are significantly different between KD and both HFD and LFD (**Figure 6F**). Together, gene expression analysis suggest a unique dysfunction of the ER-Golgi transit in KD islets which may underlie the KD-specific deficit in insulin secretion as both other groups secreted insulin in response to glucose during the clamp studies. Since upregulation in protein trafficking mainly occurs pre-Golgi, we hypothesized that a defect in Golgi processing is causing an overall defect in producing mature insulin granules and looked specifically at genes involved in Golgi organization and transport. Indeed, *Rab8b*, which was downregulated in KD islets as compared to LFD and HFD islets, plays a role in transporting proteins form the golgi to the plasma membrane suggesting reduced protein export from the Golgi.^33^ In further support of impaired Golgi functioning is our observed downregulation of *Rab30* and *Rab39* which are necessary for golgi apparatus organization.^33,34^ Finally, we compared the 175 genes that were differentially expressed in KD vs LFD and HFD vs LFD, to identify a “high-fat” transcriptomic signature. We found that HFD and KD had reduced expression of genes related to immune function (Figure S5D), suggesting the that “high-fat” signature is immune related and does not alter vesicle transport or ER/Golgi stress.

To establish the relevance of these transcriptional findings, we obtained electron microscopy images of islets from LFD and KD islets. Islets, which represent less than 1% of the surface area of the pancreas, were identified at low magnification through a flower-like structures centered around a capillary (**Figure S6A**). Β cells were identifiable by the presence of insulin granules, some of them with characteristics insulin crystals (**Figure S6A**). Insulin granules were present in β cells from both KD and LFD islets. We interrogated marked visual differences in the pool of readily releasable insulin granules at the basal membrane that could explain the lack of rapid release of insulin in KD-fed mice. However, with these granules only making up about 1% of all insulin granules and not having any known defining features,^35,36^ we could not determine whether there were differences between LFD and KD (**Figure S6A**). However, when zooming on the ER-Golgi, the Golgi apparatus in KD β cells looked vastly different than in LFD, appearing dilated and vesiculating (**Figure 6G, S6B**). This dilation, swelling, and fragmentation of the Golgi would be consistent with deficient insulin secretion observed under KD feeding.

Together, we propose a mechanism whereby a KD causes Golgi disorganization which impairs protein transport leading to compensatory upregulation of trafficking in the ER and ERGIC and a downregulation of proteins transported out of the Golgi in addition to significant ER-Golgi stress with both the stress and impaired protein trafficking suppressing insulin secretion and production (**Figure 6H**).

### 7 While a KD induces weight loss, insulin secretion becomes severely impaired

KD is currently being used to treat obesity, diabetes, and metabolic syndrome despite few preclinical studies establishing its efficacy and effects on overall metabolic health. Our observation that LT KD feeding impairs insulin secretion prompted us to evaluate insulin regulation in obese mice fed a 60% HFD subjected to KD-induced weight loss. Male and female mice rendered obese by 14 or 15-weeks of 60% HFD feeding respectively were then switched to a low-fat diet (LFD or LFMP) or a KD. Noticeably, after 15 weeks of HFD, males had reached their maximal BW whereas females weight continued to increase throughout the study (37.0g at week 15 and 55.5g at week 32, **Figure 7D**). In males, while a KD induced immediate weight loss (**Figure. 7A**), the mice on KD lost much less weight than the mice switched to LFMP and even began to gain some weight back after week 20. After 9 weeks of weight loss, the mice on LFMP had lost almost four times as much weight as the mice on KD (LFMP lost 20.9g and KD lost 5.4g) (**Figure. 7A**). Females switched to the LFD and KD rapidly and drastically lost weight for the first three weeks after starting on the new diets and then gained a small amount of weight back, though their weight still remained far below that of mice on HFD in (Figure. 7D).

**Figure 7.**
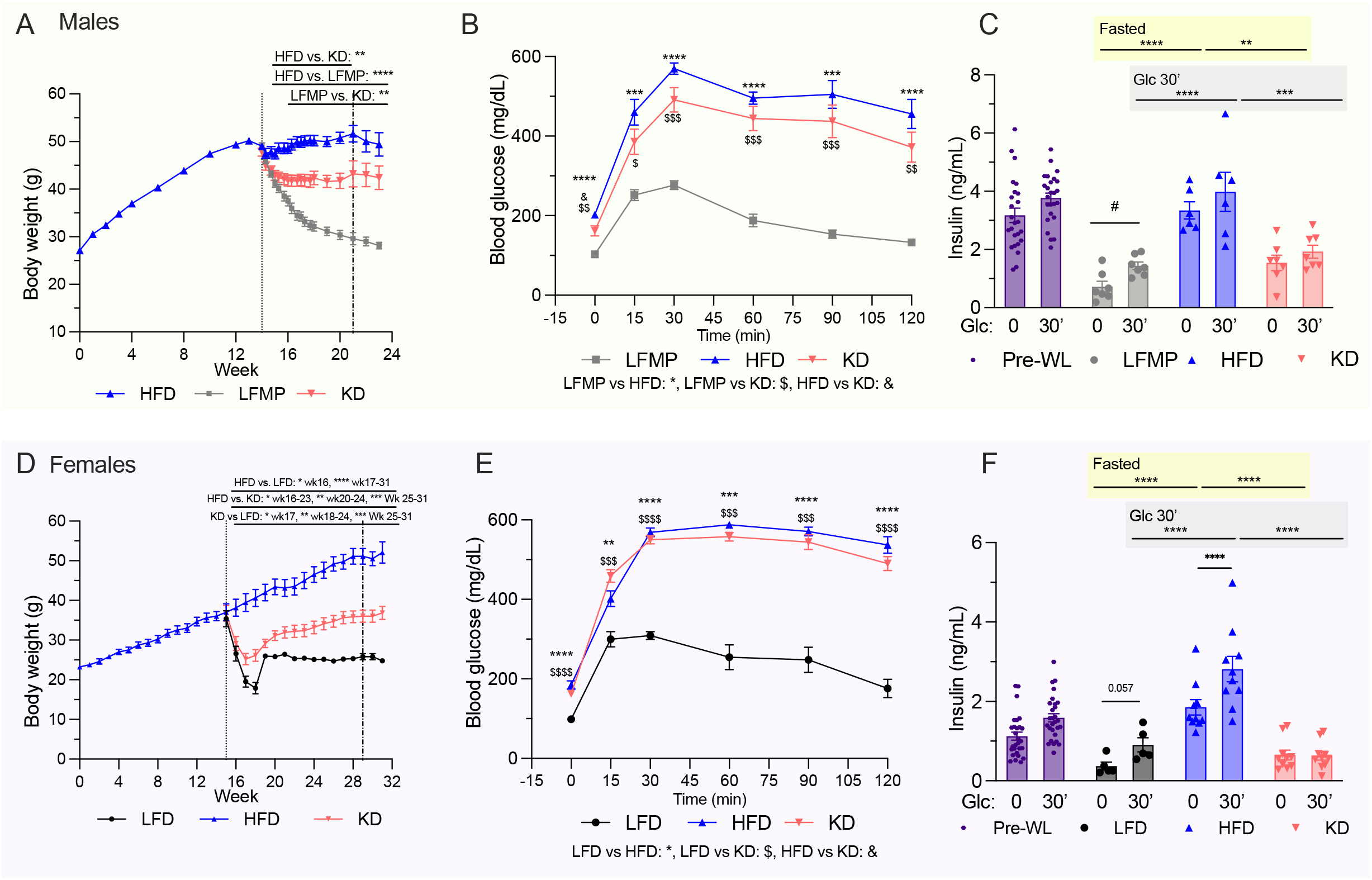
A KD for weight loss is less effective than a LFD and causes impaired insulin secretion. A-C) Weight loss intervention in male mice. A) Mice (n=30) gained weight on 60% HFD for 14 weeks before being switched to WL intervention on KD or LFMP as indicated by dotted line (n=7-10/ group) for 9 weeks. B) GTT after 7 weeks of WL (week 21 dotted line on the BW graph in A) with 1.5mg/g glucose. C) I*n vivo* GSIS: Plasma insulin during GTT in B and pre-WL. D-F) Weight loss intervention in female mice. A) Mice (n=28) gained weight on 60% HFD for 15 weeks before being switched to WL intervention on KD or LFD as indicated by dotted line (n=5-10/ group) for 16 weeks. E) GTT after 14 weeks of WL (week 29 dotted line on the BW graph in D) with 1.5mg/g glucose. F) I*n vivo* GSIS: Plasma insulin during GTT in E and pre-WL. <u>Statistics:</u> One-way or two-way ANOVA with post hoc Tukey’s HSD. Number of symbols represents significance level: * p<0.05, ** p<0.01, ***p<0.001, **** p<0.0001. Data are represented as mean ± SEM.

Like the males, the females on LFD lost much more weight than those on KD, with females on KD ending up with a higher weight than before starting the KD (LFD lost 9.7g, and KD gained 0.6g). Overall, in both males and females, KD induced weight loss, but the low-fat diets were a more effective weight loss intervention than the KD. In both males and females, the early weight loss was accompanied by an initial drastic drop in food intake when switching from HFD to either KD or LFD/ LFMP (Figure S7E, S7I). While food intake increased in mice on KD and LFD/ LFMP, the intake remained lower than that of mice that remained on HFD showing that reduced food intake can at least partially explain reductions in weight (**Figure S7E, S7F, S7I, S7J**).

Once body weight had begun to stabilize - 7 and 14 weeks into the WL interventions in males and females respectively-*in vivo* GTT and GSIS were performed. In both males and females, although the groups switched to KD had lost weight, the mice were as glucose intolerant as mice on HFD who had not lost any weight (**Figures 7B,7E**). The measurement of blood insulin levels revealed that mice switched to LFMP and LFD no longer had hyperinsulinemia and had normal GSIS and glucose tolerance response (**Figures 7B,7C,7E,7F**). In contrast, while the males and females on KD now also had low insulin levels, insulin secretion in response to glucose was gone (**Figures 7C,7F**).

These results shows that regardless of sex and regardless of body weight at initiation of the WL intervention, KD induces WL but impaired/aberrant insulin secretion occurs and prevents the normalization of glucose homeostasis.

Because a KD caused high lipid levels in our prevention study, we also measured cholesterol and triglyceride levels following weight loss to determine whether KD induced weight loss causes favorable changes in plasma lipids. We found that while LFD and LFMP induced weight loss reduced cholesterol levels, even though mice on a KD lost weight, their cholesterol and triglyceride levels remained elevated (**Figure S7G, S7H, S7K, S7L**). These findings confirm that while a KD can induce weight loss, it generally cannot reverse metabolic dysfunction.

## Discussion

The ketogenic diet has grown in popularity over the past several decades as a tool for improving weight and metabolic health. While KD benefits in treating epilepsy are concrete, its effects on metabolic health have been largely understudied. In particular, changes in glucose metabolism upon KD are not fully understood. This is of major significance since use of a KD as a nutritional approach to treat type 2 diabetes and obesity related metabolic syndrome is becoming more common and since most people who go on a KD will likely consume glucose eventually because of the difficulty to adhere long-term to the diet.^37^ In addition, the very high fat content of the diet can challenge whole body regulation of lipid homeostasis.

Our study is one of few long-term KD interventions in which mice had ad libitum access to KD. We incorporated two low fat diets, a standard LFD with 20% kcal from proteins and a LFMP with a lower 10% kcals from proteins that matches KD protein content to control for the potential confounding effects of reduced proteins. Throughout our investigations, we found that the LFD and LFMP produced virtually identical phenotypes suggesting that the effects of KD are independent of the more moderate protein content. Furthermore, other studies have shown that 10% protein is enough to support normal physiology in B6 mice.^38,39^ We also included a comparison group on 60% HFD, the gold standard model of diet-induced obesity for which metabolic disturbances are well described. Our study was powered to detect significant differences and our results were replicated in 2 to 3 independent cohorts. We performed extensive characterization of whole-body metabolic parameters, including state of the art, hyperglycemic clamp, dynamic GSIS and EM experiments. Finally, we included both sexes and carefully profiled the diet:sex interaction known to affect metabolic outcomes.^40,41^ For the most part, the phenotypic effects of KD followed the same patterns in male and female mice, yet interesting differences were observed as discussed below.

### Conflicting findings of KD on weight regulation

In our study, both male and female B6 mice fed a KD *ad libitum* gained less weight than mice on an obesogenic 60% HFD but more than mice on a LFD. A number of rodent studies have reported similar prevention of excessive weight gain on a KD as compared to a HFD^9,15,17,23^. Weight gain between KD and chow fed rodents has led to more discordant results, with several studies describing lower body weight under KD.^16,19,25,42^ Noticeably, two lifelong studies on KD, noted the necessity of restricting caloric intake^20^ or cycling the KD with a LFD^9^ to prevent obesity,^9,20^ and another 8-week study in middle-aged females also reported using pair feeding to prevent weight obesity.^43^ The long-term duration of our study may explain these discrepancies. Accordingly, in our study, it took several weeks (5 in males, 12 for females) for males and females on KD to weigh significantly more than those on LFD. Differences in diet composition, for both the KDs and LFDs/ chows, as well as specific fat composition of the KD, can all also affect the phenotypic outcomes.^7,44-46^ Future studies could directly investigate these parameters that were not explored here.

While group differences in weight were similar between males and females, the rates of weight gain were different. Males on HFD reached a BW plateau in around 13 weeks and in around 24 weeks on KD. In females, the rate of BW gain was generally lower and they continued to gain weight on HFD throughout the study while BW plateau was achieved around 28 weeks on KD. Body composition analysis revealed that males on HFD increased both fat mass and lean mass, while males on KD only significantly increased fat mass. Females body composition behaved differently in that they accrued fat and lean mass to a similar extent under HFD or KD. Differences in BW gain can be explained by differences in food intake and/or energy expenditure and KD has been described to affect both when compared to LFD [increased FI,^9,17^ increased EE^15,25,42^]. In our study, reduced caloric intake compared to HFD can at least partially explain the lower weight, although concomitant changes in energy expenditure were not assessed systematically. During our WL intervention, we observed a dramatic decrease in FI during the first couple of weeks that matches the initial drop in weight. However, weight stabilized after mice increased their *ad libitum* caloric intake to levels that were slightly below mice maintained on HFD suggesting that FI was the main driver of weight changes. Importantly, KD was less effective than chow diets at inducing weight loss as mice on LFD/ LFMP lost more than double the weight that mice on KD lost.

### KD leads to hyperlipidemia

Despite being lighter than mice fed a 60% HFD, males and females on KD had significant increases in plasma triglycerides and NEFA. KD-fed males also show hepatic steatosis and increased plasmatic ALT activity, suggestive of liver dysfunction. Those findings are in agreement with most other studies of KD.^15,17,19^ With the increasing prevalence of metabolic dysfunction associated steatohepatitis (MASH) and especially in people with obesity, the potential of KD to cause or worsen pre-existing hepatic dysfunction is of utmost concern. Surprisingly, females on KD did not display signs of hepatic lipid deposition nor liver dysfunction, and these conditions were only observed under 60% HFD. Future studies will explore these sexual dimorphisms with the potential to uncover novel biology of hepatic buffering of lipid spill over.

### KD causes severely low insulin levels

We report that severe glucose intolerance is associated with a long-term ketogenic diet. The effect of KD on glucose homeostasis in mice is controversial. Many reports initially pointed at the benefits of KD on glucose homeostasis,^15,21,28^ yet a number of recent publications report similar negative effects on glucose tolerance to what we observed.^17,25,27^ Duration of the feeding intervention may explain these discrepancies. In agreement, we show that insulin secretion is normal in the first three-four weeks of KD but becomes impaired after 4-8 week of long-term KD. We first explored insulin resistance as a determinant of glucose intolerance in KD-fed mice but ruled it out from similar ITT responses between LFD and KD-fed mice which contrasted with classically described insulin resistance in HFD-fed mice. Insights came from *in vivo* GSIS which measures insulin levels in the blood in response to a glucose bolus and revealed a lack of insulin secretion in KD-fed mice. Reviewing the literature highlighted that regardless of the positive or negative effects of KD on GTT, insulin levels in KD mice were low across the board, irrespective of diet composition, age or study duration.^15,16,42,47^ While this is often pointed to as a positive effect (i.e. prevention or reversal of hyperinsulinemia), we posit that low insulin levels on a KD are a hallmark of β cell dysfunction and can lead to other detrimental metabolic effects.

Accordingly, two prior studies described reduced β cell mass as a cause of low insulin levels.^26,27^ In our study however, extensive characterization of islets mass did not point at differences in islets mass as the underlying determinants of KD-linked insulin secretion defect. HFD-fed mice had a reduction in small islets and an increase in larger islets, in addition to increased pancreas insulin content, but no differences were observed between mice on KD and LFD. To gain further insight into the root cause of the insulin secretion phenotype in mice on KD, we performed hyperglycemic clamp studies and dynamic GSIS on isolated islets.

### KD impairs insulin secretion but not glucose sensing

Hyperglycemic clamp in freely moving animals uncovered inefficient rapid insulin secretion in KD-fed mice, a phenomenon that has not been described so far to the best of our knowledge. In mice on KD, insulin levels did not rise significantly until 90 minutes into the 120-minute study, in stark contrast to the immediate increase following the start of the glucose infusion with peak secretion at 15 minutes observed in HFD- and LFD-fed mice. Exocytosis of insulin containing secretory granules already docked at the plasma membrane^35,48^ are incriminated in the early insulin rise following glucose, suggesting that this population of insulin granules might be specifically affected by KD. To assess cell autonomous defects in insulin secretion, β cells were isolated and tested in an *ex vivo* dynamic GSIS assay. Both HFD and KD islets had reduced insulin secretion under high glucose compared to LFD islets. Upon KCl-induced depolarization, HFD islets secreted significantly more insulin suggestive of impaired glucose sensing. In contrast, islets from KD-fed mice released similarly low levels of insulin whether stimulated by glucose or KCl, indicating defective insulin secretory machinery.

Integrating *in vivo* and *ex vivo* insulin secretion results further highlight the differences in islets physiology between HFD and KD-fed mice. Despite impaired glucose sensing and secretion in isolated islets, HFD mice are hyperinsulinemic and secreted insulin in the hyperglycemic clamp. Their double amount of the largest islets and pancreatic insulin content compared to LFD and KD likely allow for this response. This was not assessed in the ex vivo GSIS as islets of different sizes were equally distributed between the groups. Islets from KD responded similarly to high glucose and KCL depolarization suggesting defective secretory capacity that is compatible with their impaired response under clamped high glucose. Yet *ex vivo,* they were still able to secrete some insulin. It is possible that the milieu further influences insulin secretory capacity. In particular, islets are in a much more lipid rich milieu (both NEFA and TG) in KD-fed mice than HFD, a parameter of which the influence is not assessed ex vivo. A number of in vitro studies have established a causal link between long term fatty acid treatment and reduced GSIS,^49-51^ however, a washout period has been shown to repair GSIS.^51^ As such, we hypothesize that removal of KD islets from their lipotoxic milieu improved insulin secretion by allowing some of the lipid to be washed out and making the islets seem comparable to HFD islets in the context of high glucose. In future studies, ex vivo GSIS performed in media that mimics *in vivo* conditions will be important for further elucidating how a KD affects cell autonomous islet function.

### KD leads to ER/Golgi stress and altered Golgi morphology causing impaired insulin granule trafficking

Transcriptomics analysis highlighted alterations in translation, protein transport, Golgi function, and ER/Golgi stress specific to KD islets. Pathways and genes associated with ER transport, COPI and II transport, and secretion from the Golgi were all elevated suggesting increased protein transport pre-Golgi. Golgi disorganization was also indicated, with transcriptional repression of Rab30 and Rab39, which are important for Golgi organization,^33,34,52^ and of Rab8b, which plays a role in Golgi vesicle secretion. From these data, we suggest a model in which Golgi related defects lead to a traffic jam of proteins in the Golgi and a compensatory upregulation of pre-Golgi transport. ER stress possibly plays a role as ER stress markers were also upregulated under KD. We speculate high-lipids in KD may play a role in the development of these cellular stresses as lipotoxicity has previously been shown to cause ER stress and disrupt ER to Golgi protein transport in islets.^53,54^ Moreover, islets from people with type 2 diabetes show an increase in ER stress markers^55^ further supporting the link between ER/Golgi stress and reduced insulin secretion. In our mice, *Creb3l3*, an ER/Golgi stress marker^56^ was the most upregulated gene in KD islets suggesting that a KD causes severe ER-golgi stress . Interestingly, CREB3 (of the same family as Creb3l3) is upregulated by palmitate in human β-cells where it attenuates palmitate toxicity and is hypothesized to act as a mediator of the Golgi stress response.^57^

Electron microscopy images of islets were obtained from an imaging expert blinded to feeding conditions and unfamiliar with our sequencing results. Electron micrographs confirmed aberrations in the Golgi which appeared dilated and vesiculating in mice on a KD. Altered Golgi morphology has been incriminated in several neurologic disorders such as Alzheimer’s Disease, Parkinson’s, and other neurodegenerative disorders.^58,59^ However, a potential role in diabetes is only emerging. A recent study characterized the disrupted protein trafficking observed in islets of diabetic LepR KO db/db mice and uncovered distended and vesiculated Golgi very similar to what we see in our KD mice.^60^ Misfolded proinsulin released from the ER is put forth as cause of Golgi dilation and altered morphology.^60^ Incidentally, misfolded proinsulin has been suggested to occur in early stages of T2D,^61^ so it is possible that increases in misfolded proinsulin cause Golgi swelling and further impair insulin trafficking and secretion.

Overall, we propose that the extra high lipids levels under KD leads to ER/ Golgi stress in islets thereby disrupting protein trafficking and causing a traffic jam of vesicles in the Golgi. With impaired protein transport and ER stress, there is insufficient readily releasable insulin available rendering the islets unable to rapidly secrete insulin in response to glucose and leading to glucose intolerance. It has been postulated that, early on, lipotoxicity causes β cell proliferation such as what is seen on HFD and, that, β cell failure, occurs in later phases of lipotoxicity.^62^ Since plasma fatty acid levels correlate with islet fatty acid levels,^63^ plasma hyperlipidemia on KD may translate to high lipid content in the islets. We propose that the extra high fat of a KD as compared to a HFD causes mice on KD to bypass the islet proliferative phase going straight to early β cell failure.

### Low insulin on KD: A benefit or a pathologic state?

Many studies have shown that a KD can lower insulin levels in humans^64-68^ and mice,^16,42^ and while this is generally taken as a sign of improved glycemic control and remission of diabetes, our results led us to question whether low insulin levels are safe and whether they actually indicate improved glycemic control. Beyond enabling glucose disposal, insulin induces the adipose tissue to uptake circulating fatty acids and inhibits lipolysis thereby reducing circulating lipids.^69-71^ During insulin resistance the ability of adipose tissue to uptake and store lipids is impaired and lipoprotein lipase becomes less responsive to insulin^72^ reducing fatty acid uptake by adipocytes. When lipid storage by adipose tissue is impaired,^73^ circulating lipids increase and ectopic lipid deposition ensue such as observed in our KD-fed mice. We hypothesize that similar effects to insulin resistance are occurring on a KD but rather than having a blunted response to insulin, the mice on KD simply do not have enough insulin to allow sufficient lipid uptake, suppression of lipolysis, or suppression of gluconeogenesis as was seen in the high fasting glucose of mice on KD. While the lack of insulin on a KD may lead to a lower body weight and less fat storage, it likely worsens metabolic health by suppressing plasma lipid clearance and benign storage of fat in adipose tissue.

### Metabolic effects of KD used for epilepsy management

KD is used to manage refractory epilepsy in children. It is rarely a lifelong intervention as confirmed by a metanalysis of 45 studies reporting that, of children put on a KD, only 45% remain on the diet after one year and 29% after 2 years.^74^ Moreover, an international consensus group statement ^2^ recommends discontinuing KD by two years with some experts recommending discontinuation by 6 months.^2^ Additionally, even while “on” KD, children often cycle on and off the KD. Metabolic disturbances have been described in epilepsy patients.

Studies around the world have reported higher than average occurrence of diabetes in people with epilepsy.^75-77^ The reasons for this co-occurrence are unclear ^78,79^ but one hypothesis is that epilepsy treatments increase the likelihood of diabetes.^80^ High plasma lipids have also been reported in some patients on KD.^13,81,82^ Pancreatitis, for which hypertriglyceridemia is a risk factor, and cardiac complications have also been reported.^2,83,84^ A study of epilepsy patients on KD for at least six years concluded that the patients were at higher risk of other health complications.^82^ Overall, metabolic complications are reported in people who use KD as epilepsy treatment, but since KD is predominantly used in children and discontinued after a short time, long term studies of metabolic effects are generally lacking and warranted for translation of this dietary intervention to metabolic disease management.

In Summary, while a KD can prevent and treat obesity, it causes hyperlipidemia, hepatic steatosis, and glucose intolerance. Unlike mice on HFD, the mice on KD do not have detectable insulin resistance or hyperinsulinemia. Instead, they have impairments in insulin secretion due to a blockade in protein trafficking resulting from dilation of the Golgi apparatus which also causes ER/Golgi stress.

### Conclusions and Limitations

Our study shows that while C57Bl/6J male and female mice on KD are protected from weight gain as compared to mice on a traditional HFD, the mice experience severe glucose intolerance, high plasma lipids, and impaired insulin secretion with males also developing hepatic steatosis. Further studies in other strains of mice, other animal models, and in humans are necessary to determine whether KD-linked metabolic derangements are universal. It has been suggested that saturated versus unsaturated fats and different macronutrient compositions can have different effects. Future studies to elucidate whether the type of fat and composition of the KD influences the metabolic outcomes are important for making KD safer for all including those who require a KD to treat their epilepsy. Despite these limitations, our findings have relevant translational ramifications. They suggest that a KD used as a long-term dietary intervention may have harmful effects on metabolic health, especially in terms of β cell function, plasma lipids, and liver health, and caution its systematic use as a health promoting dietary intervention.

## Acknowledgements

This work was supported by funds from NIA R01AG065993 to AC and NIDDK F32DK137475, T32DK110966 to MRG. WLH receives funds from DK108833, DK112826 and HL170575. K.J.E. is funded in part by Damon Runyon-Rachleff Innovation Award DR 61-20, NCI R01CA222570 and the American Cancer Society (RSG-22-014-01-CCB). We thank additional members of the Chaix lab, Elena Gonzalez Alvarado, Ceyda Ural, Navya Murahari for assisting with WL study; Joseph Varre and Daniela Ramos from Holland lab for help preparing islets and performing the surgeries. We acknowledge the Metabolic Phenotyping Core (CLAMS and Bruker Minispec NMR; Dr. Ying Li), Cell Imaging Core (Axioscan Slide Scanner; Dr. Anton Classen, and Bill James), and the Electron Microscopy Core at the University of Utah for use of equipment and thank the staff for their assistance. Research reported in this publication also utilized the High Throughput Genomics Core (Illumina Nextgen Sequencing; Dr. Brian Dalley) and the Bioinformatics Core (RNAseq analysis; Dr. Chris Stubben) at Huntsman Cancer Institute at the University of Utah supported by NCI P30CA042014. *Ex vivo* GSIS was performed by the Islet and Pancreas Analysis (IPA) Core supported by the Vanderbilt Diabetes Research Center DK20593.

## Author contribution

Conceptualization, MRG and AC; Methodology, MRG, AC, WLH, WL; Investigation, MRG, RFLV, ETM, PDM, FES, SR, WL, KJE; Writing – Original Draft, MRG; Writing –Review & Editing, MRG and AC; Funding Acquisition, AC; Resources, AC, WLH, KJE, WL; Supervision, AC.

## Methods

### Mice

C57Bl6/J male and female mice from Jax (Cat#000664) were used in all studies. After receiving mice from Jax, they were placed on chow and allowed to acclimate for at least 3 weeks. When mice were 14-15 weeks old, they were placed on their respective diets. Mice had 24 hour *ad libitum* access to food and water and were on a 12:12 light:dark cycle. Lights on a 7:00am. Th prevention studies (figures 1-6) used169 males from three independent cohorts and 60 females from two independent cohorts. The weight loss studies (figure 7) used 30 female and 30 male mice. The male weight loss study was repeated in two additional cohorts (data not shown).

### Diets

All diets used in the studies were ordered from Research Diets. LFD: 10% kcals from fat, 70% kcals from carbohydrates, 20% kcals from protein (D12450K). LFMP: 10% kcals from fat, 80% kcals from carbohydrates, 10% kcals from protein (D21110806). HFD: 60% kcals from fat, 20% kcals carb, 20% kcals protein (D1292). KD: 89.9% kcals from fat, 0.1% kcals from carbohydrates, 10% kcals from protein (D16062902).

<u>Body composition</u> was measured using a Bruker Minispec small animal NMR (Bruker, Cat#LF50). Animals were awake and freely moving. Measurements were taken during the dark cycles and animals were in the ad libitum fed state.

### Blood glucose and blood ketone beta-hydroxybutyrate

were collected using handheld meters. Venous blood was obtained via a small nick in the tail. Blood glucose meters (Amazon, Nova Max Plus) had a range of 20mg/dl-600mg/dl, so any readings that were high were given a numeric value of 600. Ketone body meters (Amazon, KetoneBM) measured only the presence of beta-hydroxybutyrate.

### GTT and ITT

Before studies, mice were fasted over night for roughly 15 hours. Blood glucose levels were measured at baseline and then at regular intervals during the study as noted. Mice were given IP injections of 1.5mg/g glucose (Sigma Aldrich, D-(+)-Glucose, Cat# G7021) or .75IU/ kg insulin (Lily Medical, Humulin R U-100) in phosphate buffered saline (PBS). Blood glucose and blood was collected as noted in the experiment. For in vivo GSIS, blood was collected using a cheek bleed before and during the GTT.

### Plasma metabolites

All blood metabolite measures were performed using plasma collected from the submandibular vein. Fed blood was collected in the ad libitum fed state during the middle of the dark cycle (12pm), and fasted blood, which was taken to measures baseline insulin during in vivo GSIS was collected at 7am (onset of the dark cycle) following a 14 hour overnight fast. Blood was collected in EDTA tubes (Sarsdtet,16.444.100) and placed immediately on ice. Blood was then spun at 1000g for 10 minutes. Cholesterol, triglycerides, and non-esterified fatty acids were measured using colorimetric kits (Sekisui Diagnostics, TG: Cat#236-60, Chol: Cat#234-60, NEFA: Cat#s:999-34691, 991-34891, 995-34791, 993-35191, 997-76491). Insulin was measured using an insulin ELISA (Crystal Chem, Cat#90080).

### Liver triglycerides

Liver was collected at sacrifices and immediately snap frozen. Tissues were homogenized using a mortar and pestle kept cold with dry ice and liquid nitrogen. Lipids were extracted using a modified Folch methods. Liver (50mg) was extracted in 2:1 Chloroform: Methanol mixture (Chloroform: Sigma Aldrich, C2432-1L; Methanol. Mixture was vortexed and left at 60 C for 30-60 minutes. The aqueous layer was transferred to a new tube and 125ul water was added for every 500ul chloroform/methanol mixture. The mixture was vortexed and then spun at 500g for 10 minutes to cause phase separation. The lower organic layer was moved to a new and liquid was evaporated. Chloroform was added back to resuspend the lipids (20:1 ratio original tissue weight to chloroform). 100ul of chloroform mixture was combined with 200ul 1% triton-100 and re-evaporated at 45C. The leftover lipid triton mix was resuspended in water. Triglycerides were measured using the same triglyceride kit as for the plasma (Sekisui Diagnostics, Cat#236-60).

### Hyperglycemic clamp

#### Surgery

Mice were anesthetized using isoflurane (Vet OneNDC,13985-528-60; Kent Scientific Somnosuite, 22-01), given anesthetics, and skin was shaved and prepped with betadine and isopropyl alcohol. A small incision was made starting from halfway between the left armpit and midline and extended up roughly 50cm to the next. The jugular vein was isolated and the catheter (Instech, C20PU-MJV2013) was introduced into 1-2mm incision in the jugular. The catheter was slid roughly 10mm into the vein, and ligatures (Fine Science Tools, Cat#18030-60) were applied above and below catheter insertion to hold it in place. The catheter was then threaded around the side of the mice and pulled through a small incision sitting atop the scapula. Incisions were sutured closed (Covidien, SN1955), The catheter was held in place using a backpack harness (Instech, Cat#CIH62) and filled with lock solution. Mice were allowed to recover for four days.

#### Clamp Procedure

On the day of the study catheters were connected to a swivel system (Instech, Cat#KVAH62T) system allowing mice to be freely moving during the clamp. Blood glucose was measured beginning 90 minutes before the clamp to ensure mice had a steady baseline glucose. Mice were infused with 50% glucose in saline starting at a rate of 500ul/ hour for 2 minutes. After the first two minutes, the infusion rate was changed to an amount in microliters that was 3x the body weight (is a 50g mouse received an infusion of 150ul/hr). Then the infusion amounts were changed based on blood glucose levels that were taken every 3-5 minutes with a target glucose level of 250mg/ dl. Tail bleeds were also performed at baseline 0, 5, 10, 15, 20, 25, 30, 60, 90, 120 so that insulin levels could be measured.

#### Islet Harvest for RNA sequencing

Mouse was sacrificed and the pancreas was perfused using an oxygenated buffer of HBSS (Thermofisher Scientific, Cat#14025092) with 25mM HEPES (Gibco, Cat#15630-080), 8mM glucose, and 0.2% BSA. 0.1mg/ml Liberase (Sigma Aldrich, 0540102001) was added tto the buffer for pancreas digestions. The pancreas was digested at 37C for 30 minutes. After 30 minutes, the samples were flooded with fresh buffer and allowed to settle for 10 minutes so that the islets would sink to the bottom and the debris would float to the top; part of the supernatant was removed, fresh buffer was added to replace the volume removed, and samples were agitated to mix. This process was repeated until the liquid was very clear (4-6 times); the islets with the clear liquid were then handpicked under a microscope. Islets were then pelleted, washed with PBS, and snap frozen.

#### RNA sequencing and analysis

RNA was extracted using Trizol (Invitrogen, Cat#15596018) Chloroform and followed by a Qiagen RNEasy kit with DNASe treatment (Islets: micro kit 74004, Liver: Mini Kit 74106) . The University of Utah High Throughput Genomics core performed RNA sequencing with the following methods. Total RNA samples (5-500 ng) were hybridized with NEBNext rRNA Depletion Kit v2 (Human, Mouse, Rat) (E7400) to substantially diminish rRNA from the samples. Stranded RNA sequencing libraries were prepared as described using the NEBNext Ultra II Directional RNA Library Prep Kit for Illumina (E7760L). Purified libraries were qualified on an Agilent Technologies 4150 TapeStation using a D1000 ScreenTape assay (cat# 5067-5582 and 5067-5583). The molarity of adapter-modified molecules was defined by quantitative PCR using the Kapa Biosystems Kapa Library Quant Kit (cat#KK4824). Individual libraries were normalized and pooled in preparation for Illumina sequence analysis. *The following adapter reads were used: Adapter Read*1: AGATCGGAAGAGCACACGTCTGAACTCCAGTCA; *Adapter Read 2*: AGATCGGAAGAGCGTCGTGTAGGGAAAGAGTGT. Sequencing libraries were chemically denatured in preparation for sequencing. Following transfer of the denatured samples to an Illumina NovaSeq X instrument, a 150 x 150 cycle paired end sequence run was performed using a NovaSeq X Series 10B Reagent Kit (20085594).

Data were aligned to mm39 and deseq2 analysis was run as described by Love et. al.^86^ by the University of Utah Bioninformatics Core. Pathway analysis was performed using Reactome^87^ Camera analysis on differentially expressed genes. Metascape^88^ was also run on differentially expressed genes to identify gene ontology groups that were enriched. UpSet^89^ plots were used for data visualization.

Fastq files are available online. Liver RNAseq: GEO, GSE244492; Islet RNAseq: GE, GSE268020

#### Insulin staining and islet sizing

Pancreas was collected after 29 weeks on the respective diets fro both male and female mice. In males, 8-12 slices were taken throughout the entire pancreas from n=4-6 animals per groups for a total 5157 islets analyzed on LFD, 6718 on HFD, and 6389 on KD. For females, 6-7 slices were taken throughout the entire pancreas (n=5/ group), for a total of 3749 LFMP and 3428 KD islets analyzed.

Sections for histology were collected at sacrifice and then placed in 10% formaldehyde (Ricca, Cat#3190-5) for 24 hours followed by 70% ethanol (Decon Laboratories, Cat#2701). Sections were then paraffin embedded and sectioned to 5um thickness by the University of Utah Research Histology Core. Sections were deparrafinized and rehydrated in sequential 3-minute soaks in xylene (x2) (Sigma Aldrich, Cat#296325) 100% ethanol (Decon, Laboratories, Cat# 2701), 95% ethanol, 70% ethanol, 50% ethanol, and water. Antigen retrieval was performed using 10mM Sodium Citrate (Fisher Scientific, BP327-500) with 0.05% Tween 20 (Calbiochem, 655204) pH6 in kept around 95C for 30 minutes. A 30-60 minute blocking step with 2% bovine serum albumin (BSA) (Sigma Aldrich, Cat#A7906) in tris buffered saline was performed, followed by 30% incubation with 3% hydrogen peroxide to quench endogenous peroxidase activity. After washing 3x for five minutes with PBS + 0.05% Tween 20 the HRP conjugated insulin antibody was used (Abcam Cat#28063 mouse anti insulin and proinsulin-AB dilution 1:200) in the dark for 60 minutes. Antibody was washed off as described above and followed DAB application (Thermo Scientific, Cat#34002) and a hematoxylin counterstain (Vector Labs, Cat# H-3502). Slides were then dehydrated and coverslipped.

Slides were imaged on an Axioscan 7 slide scanner (Zeiss), and images were processed in Qupath^90^. To quantify insulin positive area, thresholding for DAB staining was performed, and manual corrections were made as needed to remove artefacts or add islets that were not counted. A minimum islet size was set to exclude islets with an area of less than 100um^2^ this corresponds to a diameter of roughly 11.28um which is slightly larger than a singular beta cell, and detections smaller than 100um^2^ often did not contain a visible nucleus. We calculated area and perimeter for each unique islet. A separate analysis in Qupath for total pancreas area was created. We created a positive area detection threshold for hematoxylin and measured the total area; regions of adipose tissue surrounding the pancreas were removed. Area and perimeter data for islets and the pancreas sections were extracted from Qupath.

The binned areas used in the frequency distribution correspond to islet diameter as follows: 100μm^2^= diameter of 11.3μm or the size one beta cell (and the minimum size needed to identify a nucleus and clear borders in the analysis), 170μ^2^= diameter of 14.7μm, 490.87μm^2^= diameter of 25μm, 1963μm^2^= diameter of 50μm, 4417.86μ^2^= diameter of 75μm, 7853.981μm^2^= diameter of 100μm, 17671μm^2^= diameter of 150μm.

#### Liver histology and scoring

Sections for histology were collected at sacrifice and then placed in 10% formaldehyde (Ricca, Cat#3190-5) for 24 hours followed by 70% ethanol (Decon Laboratories, Cat#2701). Sections were then paraffin embedded, sectioned to 5um thickness, and stained with hematoxylin and eosin (H&E) by the University of Utah Research Histology Core. Slides contained one section from each diet group and were scored by a board-certified pathologist (K.J.E.) who was blinded to the diet conditions. Steatosis was determined by estimating percent area containing lipid droplets, and liver inflammation was scored between 0-3 based on the scale developed by Liang et al.^85^ Images were acquired using cellSens software and an Olympus DP73 camera attached to an Olympus BX53 microscope with UPlanFL objectives.

#### Electron Microscopy

Mice were anesthetized by isoflurane gas Mice were anesthetized using isoflurane (Vet One, NDC,13985-528-60; Kent Scientific Somnosuite, 22-01), and fixed by cardiac perfusion with 2.5% glutaraldehyde and 1% paraformaldehyde, 6mM CaCl2, and 2.4% sucrose in 0.1M cacodylate buffer, pH 7.4 (Electron Microscopy Sciences, CAT#16537-07).

Immediately after perfusion, the pancreas was dissected out and soaked in fresh fixative, where the organ was sliced into small blocks of about 10 mm^3^. Tissue blocks were transferred into fresh fixative where they were continuously fixed for at least two hours at room temperature before being stored at 4°C.

Sample processing and conventional electron microscopy were conducted by the University of Utah Electron Microscopy Core Facility using a microwave assisted-fixation and processing protocol (Pelco Biowave®, 36500). Sequential steps were as follows: 2% osmium post-fixation, 4% uranyl acetate en-bloc staining, serial dehydration in ethanol/acetone, and embedding in Embed 812 (Electron Microscopy Sciences, Cat# #14120). Specimens in the cured epoxy-resin block was sectioned using Lecia ultramicrotome UC6. Islet of Langerhans was identified under a light microscope in semithin sections stained with toluidine, then ultrathin sections of 100 nm thickness were cut and deposited onto carbon-formvar coated slot grids. Sections on grids were further contrast-stained with uranyl acetate and lead citrate and examined at 120KV with JOEL JEM 1400 equipped with a GatanOrius digital camera.

#### Whole pancreas insulin quantification

At sacrifice pancreas was carefully dissected and weighed. A small piece (10-20mg) was taken and weighed separately. This small piece was added to a tube containing 5ml of acid ethanol extraction buffer (1.5ml 1N HCl with 98.5ml 70% ethanol). The pancreas was homogenized by vortexing with metal screws for 2 hours on two consecutive days or until mixture was homogenized. The slurry was then centrifuged at 2000g for 3 minutes to pellet any debris. 500ml of supernatant were mixed with 500ml 1M Tris. The Tris extract mixture was then diluted 1:100 using Crystal Chem Islet sample diluent and then run on the Crystal Chem ELISA using the high range assay.

#### Ex vivo GSIS

The ex vivo GSIS was performed by the Vanderbilt Islet and Pancreas Core using established protocol that are found online: Islet Harvest^91^; Islet Perifusion ^92^.

#### Islet harvest

Mice were anesthetized with ketamine/xylazine solution (1 μl/mg body weight) and after loss of consciousness they were placed on a surgical stage and skin was prepped with 70% ethanol. The skin and peritoneum were opened, and the bile duct was located and closed with a suture at the duodenum. Using a bent 30G needle collagenase solution (0.6 mg/mL collagenase P in HBSS) (Collagenase: Millipore Sigma Cat#11249002001; HBSS: Thermofisher Cat# Cat#14025) was injected into the bile duct entering the duct near the liver to inflate the pancreas. The pancreas was removed and placed in a conical tube containing 5ml HBSS/ Collagenase solution, the tubes were placed in a 37C water bath shaker for 8 minutes with shaking by hand every 4 minutes and until the tissue had dissolved into a slurry at the end of the second shaking.

Tubes were placed on ice and topped up to 15mls with HBSS with 10% FBS (Sigma Aldrich, Cat#F0926) and then centrifuged at 200 rcf for 00:01:00 at 25 °C. Liquid was decanted, and pellets were washed by adding 10 mL HBSS with 10% FBS and then vortexed to mix before centrifuging again. This process was repeated 3x. After the final wash, supernatant was discarded, and 10 mL HBSS with 10% FBS was added and mixed. The slurry was transferred to a 50 mL conical tube. The original tube was washed with another 5 mL HBSS with 10% FBS and added to the 50 mL tube for a total of 15 mL of digest. The digest was vortexed, 15 mL of Histopaque gradient was underlayed using a manual bulb pipette. Tubes were centrifuged at 500 rcf for 00:10:00 at 25 °C. New 50 mL tubes containing 15 mL HBSS with 1% FBS were prepared, and islet layer was transferred to the new tubes. Tubes were centrifuged tubes at 500 rcf for 00:10:00 at 25 °C, the liquid was decanted and pellets were washed 3X with 25 mL HBSS with 1% FBS vortexing each time after adding HBSS to mix before centrifugation. After the final decanting, 10 mL HBSS with 10% FBS to were added to the tube, and mixed by pipetting. The mixture was transferred to a 10 cm untreated culture dish. Islets were handpicked under an inverted microscope at 4x. Islets were then sized and handpicked to a 6cm dish.

#### Perifusion

Base perifusion media: 1L DI water plus the following: 3.2 g NaHCO3 (Sigma Aldrich, Catalog #S6014-500G), 0.58 g L-Glutamine (Sigma Aldrich, Catalog #G8540-100G), 0.11 g Sodium Pyruvate (Sigma Aldrich, Catalog #P2256-25G), 1.11 g HEPES (Sigma Aldrich, Catalog #H7523), (8.28 g DMEM (Sigma Aldrich, Catalog #90113), 1 g RIA-grade BSA (Sigma Aldrich, Catalog #A7888), and 70 mg L-ascorbic acid (Sigma Aldrich, Catalog #A5960).

5.6mM glucose media was made by combining 550ml base media with 0.5549 g Glucose (Sigma Aldrsich, Catalog #D16). 16.7 mM high glucose media was made byby adding 0.7522 g Glucose (Sigma Aldrich, Catalog #D16) to 250 mL base perifusion media. KCl media was mice by by adding 0.149 g Potassium chloride (Sigma Aldrich, Catalog #BP366-500) to 100 mL 5.6 mM glucose media.

Perifusion setup was made up of a circulating water bath, a fraction collector, a peristaltic pumps capable of pumping 1 mL/min, and glass columns with 2 fixed-end pieces (Diba Omnifit® LC columns (Cole Parmer, Catalog #006CC-10-05-FF) as described^92^.

Islets were transferred to a clear 1.5-mL microcentrifuge tube and then loaded into perifusion system. Bubbles were removed, before starting the system, and fraction collection was set to every 3 minutes.

After collection of 10 preliminary fractions using 5.6mM glucose baseline media to wash the islets, the experiment started. Baseline media was used from 0-12 minutes, 16.7mM glucose media was used from 12-42 minutes, and then intake was switched back to baseline media from 42-63 minutes. Next, 5.6mM glucose with 20mM KCl was used from 63-72 minutes, and then the media was switched back to the baseline media for the rest of the experiment.

Islets were recovered to a 6-cm dish to calculate IEQ after the perifusion.

##### QUANTIFICATION AND STATISTICAL ANALYSIS

Graphpad Prism was used for all statistical analyses. A P value of ≤ 0.05 was used as the cutoff for significance. All results and tests used can be found in the relevant figure and figure legends. Graphs show all individual data points with the bar showing the mean and the error bars showing standard error of the mean. One-way or two-way ANOVA were used for all group comparisons with post-hoc Tukey’s HSD test aside from sequencing which used Deseq2 analysis and a Benjamini-Hochberg correction for the adjusted P-value. Sample sizes were determined with a power calculation, and animals were randomly assigned to groups that were matched for body weight and spread.

