## Supplemental Figures & Legends for "Long-term ketogenic diet causes hyperlipidemia, liver dysfunction, and glucose intolerance from impaired insulin trafficking and secretion in mice"

**Figure S1 related to Figure 1.**
A) Respiratory exchange ratio (RER) measured between 13-17 weeks on diets (n=4/ group).
B-E) Body composition by percent of BW in males corresponding to figures 1F and 1G (B: % fat, D: % lean; n=4-6/group), and females corresponding to figures 1J and 1K (C: % fat, E: % lean; n=9-10/group).
F-G) Food intake over time in males (F) (n=8-14 cages) and females (G) (n=2-4 cages)
Statistics: Mixed effects ANOVA, post Hoc testing used Tukey's HSD test. * $p<0.05$, ** $p<0.01$, ***$p<0.001$, **** $p<0.0001$. Data are represented as mean ± SEM.

**Figure S2 related to Figure 2. Mice on KD and HFD develop high cholesterol.**
A, B) Plasma cholesterol levels in (A) male mice, 32 weeks on diets, n=5-10/ group and (B) female mice, 15 weeks on diets, n=9-10/ group. All blood samples were collected in the fed state in the middle of the dark phase.
Statistics:
A, B) One-way ANOVA with Tukey's post hoc testing with p values as: * $p<0.05$, ** $p<0.01$, ***$p<0.001$, **** $p<0.0001$. Data are represented as mean ± SEM.

**Figure S3 related to Figure 3. Glucose intolerance in mice on KD is independent of delivery route and insulin secretion defects develop between 3- and 8-weeks on KD**
A, B) Oral glucose tolerance testing using a bolus of 1.5mg glucose/ g body weight. A) males after 20-24 weeks combined from two independent cohorts (n=14-16/ group). B) females at 17 weeks on diets (n=8-10/group)
C) Meal tolerance testing using Ensure Plus (10ul/g BW) males at 25 weeks n=8-10/ group.
D-F) ipGTT males n=5-group with a bolus of 1.5mg/g given intraperitoneally.
G-I) Plasma insulin levels before and 30 minutes after a 1.5mg/g glucose bolus.
D, G) After 7 days on diets (n=5/ group).
H, I) After 3 weeks on diet (n=5/ group).
F, I) After 8 weeks on diets (n=5-/10/ group).
Statistics: Two-way ANOVA with Tukey's HSD test: number of symbols represent * $p<0.05$, ** $p<0.01$, ***$p<0.001$, **** $p<0.0001$. Data are represented as mean ± SEM.

**Figure S4 related to Figure 4. A KD does not increase islet size or pancreas insulin content.**
A, B) Analysis of chromogenic insulin/ proinsulin staining in the pancreas with hematoxylin counterstain. Qupath was used to quantify islet area based on detection of the brown insulin stain. 8-12 slices were taken throughout the pancreas from n=4-6 animals per groups for a total 5157 islets analyzed on LFD, 6718 on HFD, and 6389 on KD.
A) Percent insulin positive area (total insulin positive area/ whole slice area*100)
B) Frequency distribution of islets sizes as counts in each bin per slice.
Bins correspond to islet diameter $100\mu m^2$= diameter of $11.3\mu m$ or the size one beta cell (and the minimum size needed to identify a nucleus and clear borders in the analysis), $170\mu^2$= diameter of $14.7\mu m$ , $490.87\mu m^2$= diameter of $25\mu m$, $1963\mu m^2$= diameter of $50\mu m$ , $4417.86\mu^2$= diameter of $75\mu m$, $7853.981\mu m^2$= diameter of $100\mu m$, $17671\mu m^2$= diameter of $150\mu m$.
C) Male total pancreas insulin content after 6 months on the diets n=5-6/ group.
D) Female pancreas insulin content after 49 weeks on respective diets n=3-4/ group.
Statistics: One way or 2 way ANOVA post hoc Tukey's HSD for multiple comparisons.
* $p<0.05$, ** $p<0.01$, ***$p<0.001$, **** $p<0.0001$. Data are represented as mean ± SEM.

**Figure S5 related to Figure 6.**
Bulk RNA sequencing from islets isolated from male mice after 36 weeks on interventions analyzed using Deseq2.
A-B) Volcano plot of all genes between specified comparisons. Orange indicates significantly lower expression, and purple indicates significantly increased expression.
C-D) Differentially expressed in the indicated comparison were run through Metascape which identified ontology categories that were enriched in one group versus the other as displayed in the heatmap; the purple indicates increased expression in the corresponding group, and the color intensity represents -log10(P-value) of the enrichment as calculated by Metascape.
Statistics: Differentially expressed genes were identified using DESeq2 analysis with a Wald test and Benjamini-Hochberg correction. Significant genes had an adjusted P-value of <0.05.

**Figure S6 related to Figure 6.**
A) Electron micrographs showing identification of islets and zooming in on the islets and beta cells.
B) Electron micrographs showing more views of the golgi from LFD and KD beta cells.

**Figure S7 related to Figure 7.**
A, B) Males after 13 weeks of weight gain. (A) GTT with 1.5mg/g glucose and B) I*n vivo* GSIS taken during GTT; n=25.
C, D) Females after 14 weeks of weight gain. (C) GTT with 1.5mg/g glucose and D) I*n vivo* GSIS taken during GTT; n=28.
E) Food intake over time and (F) average food intake during weight gain or weight loss in males (n=2-6 cages).
G) Plasma cholesterol and H) triglyceride levels after 7 weeks of weight loss measured in the refed state during the middle of the dark phase (n=7/ group).
I) Food intake over time and (J) average food intake during weight gain or weight loss. in females (n=2-6 cages).
K) Plasma cholesterol and L) triglyceride levels after 14 weeks of weight loss measured in the middle of the dark cycle 3 hours after food removal.
Statistics: Paired T-test used in B and D, otherwise one-way or two-way ANOVA with post-hoc Tukey's HSD were used; * p<0.05, ** p<0.01, ***p<0.001, **** p<0.0001. Data are represented as mean ± SEM.

Figure S1 related to figure 1

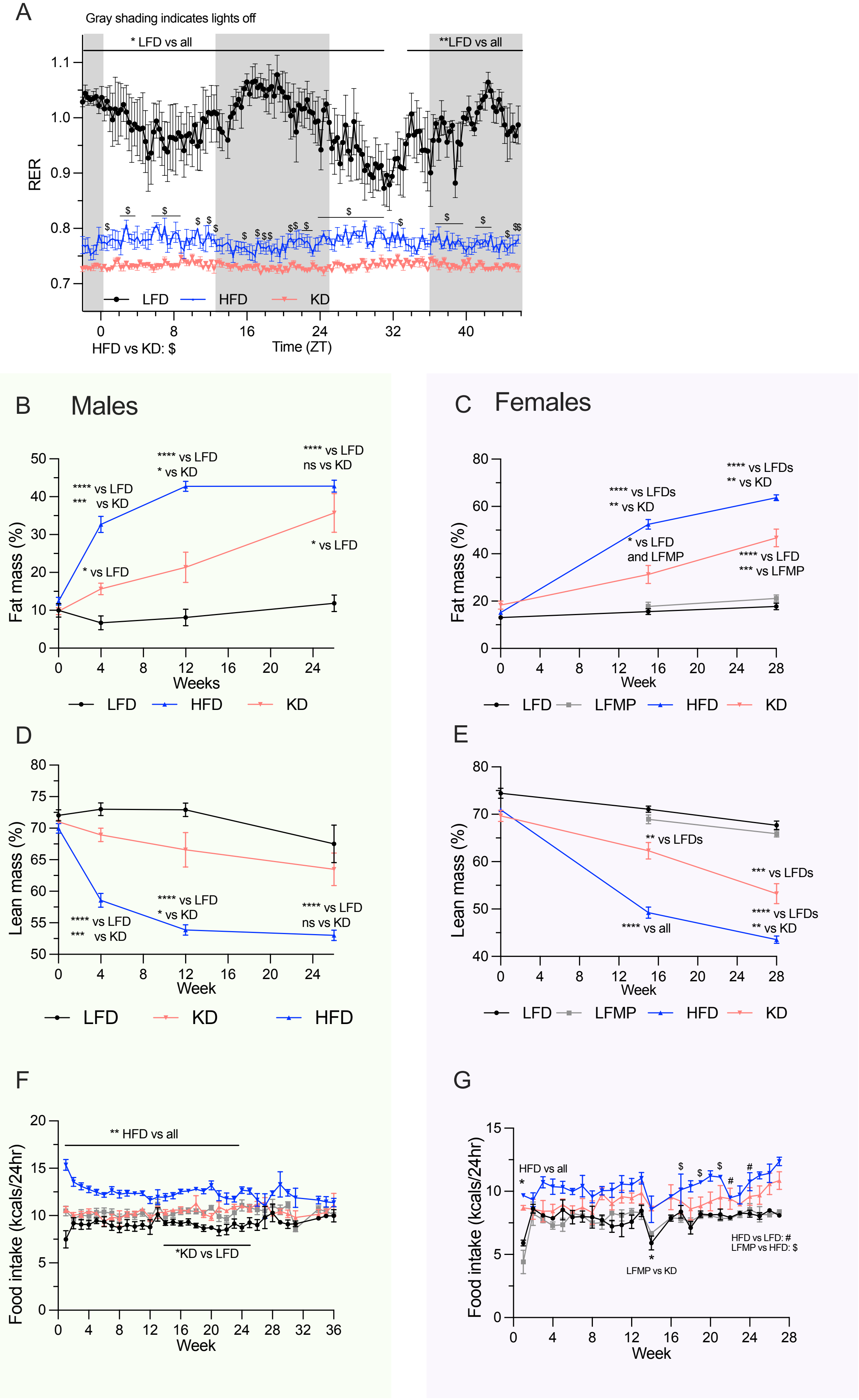

Figure S2 related to figure 2

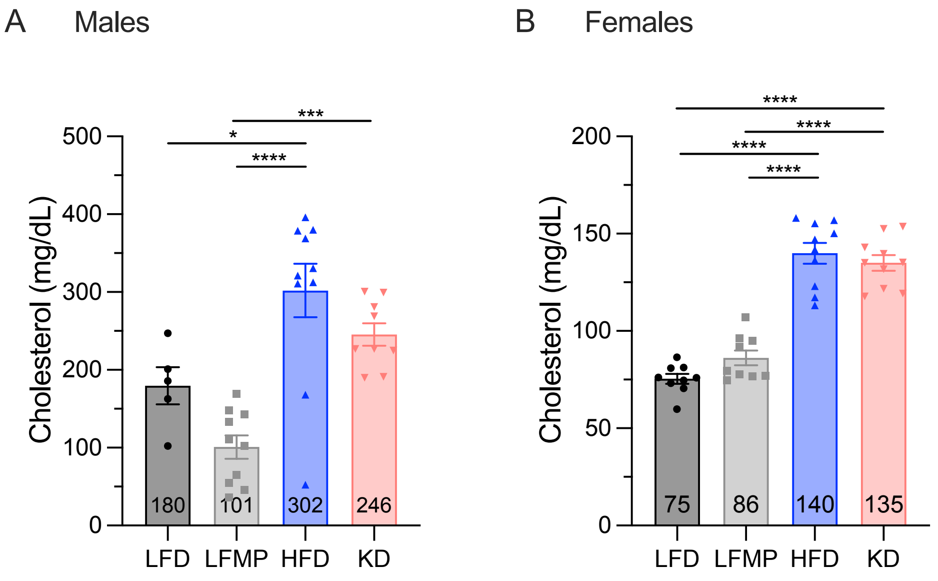

Figure S3 related to figure 3

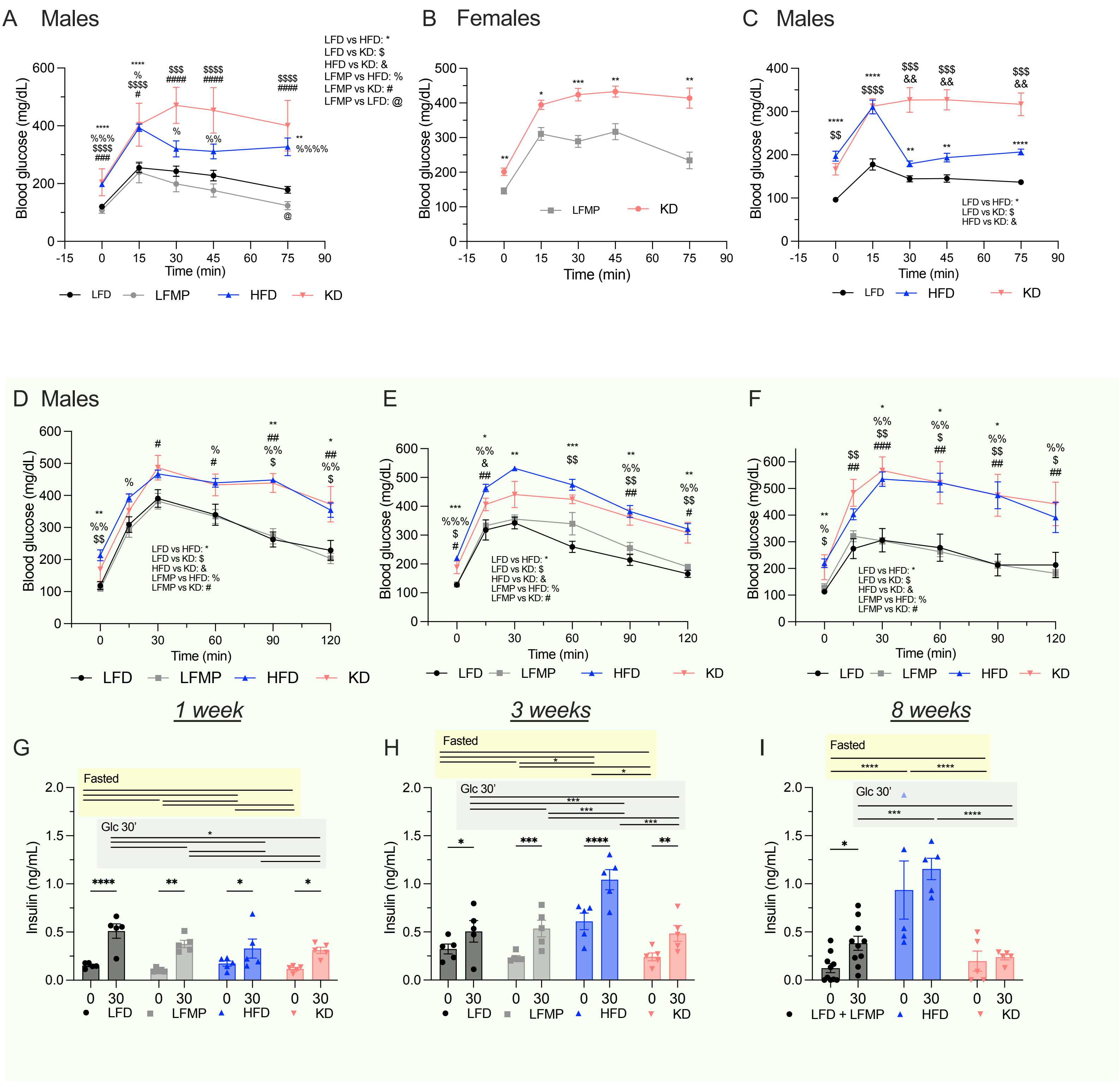

Figure S4 related to figure 4

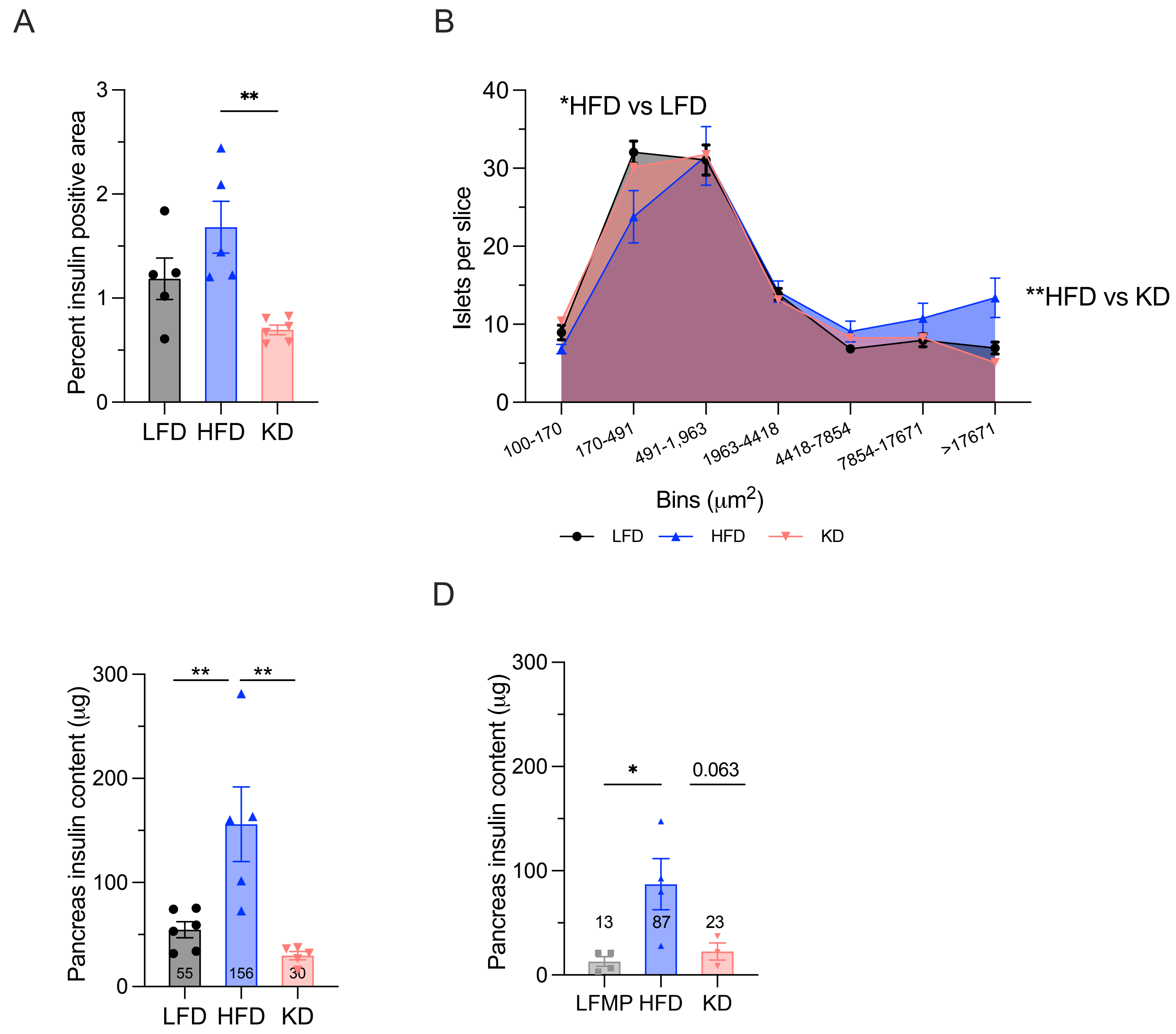

Figure S5 related to figure 6

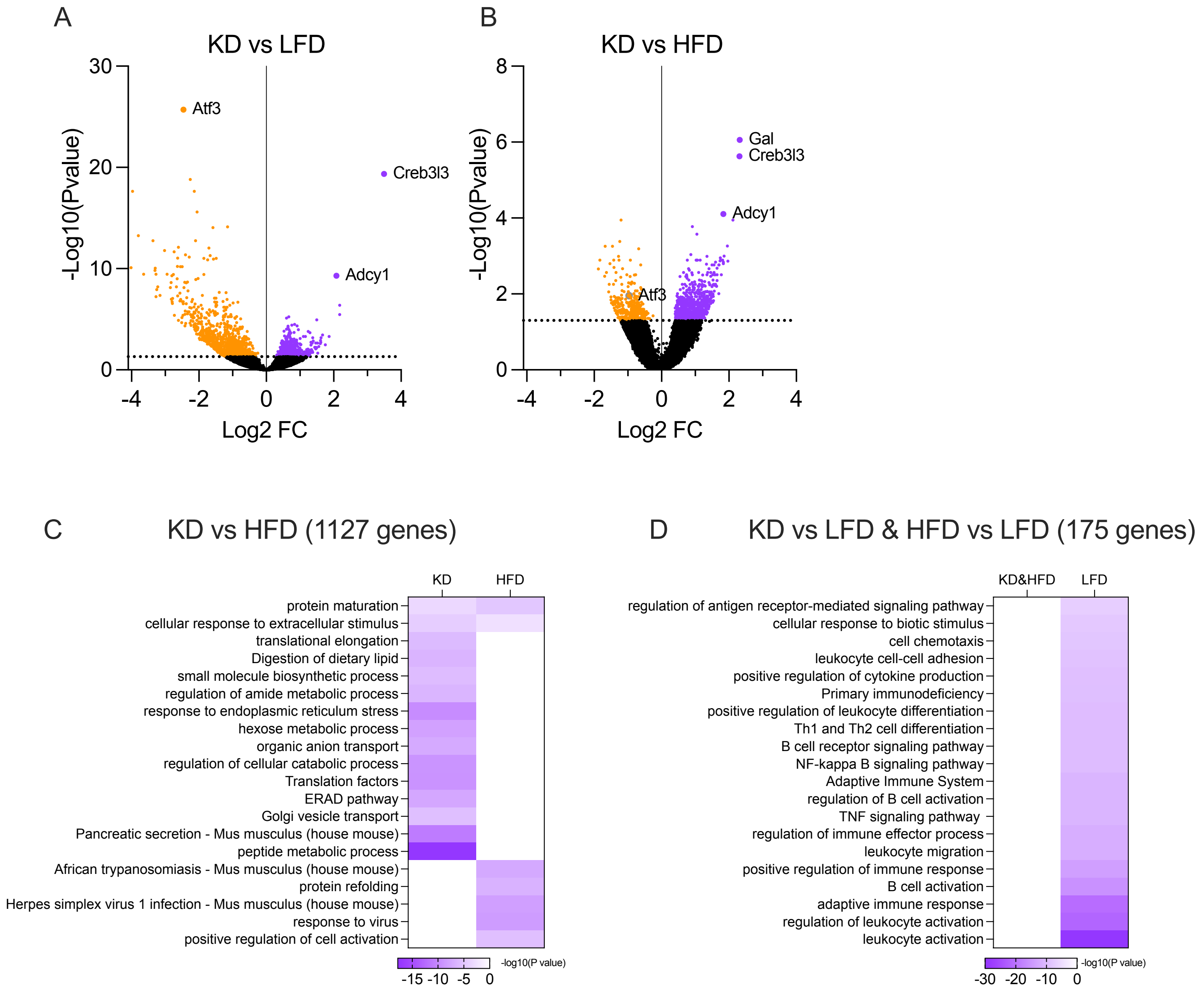

Figure S6 related to figure 6

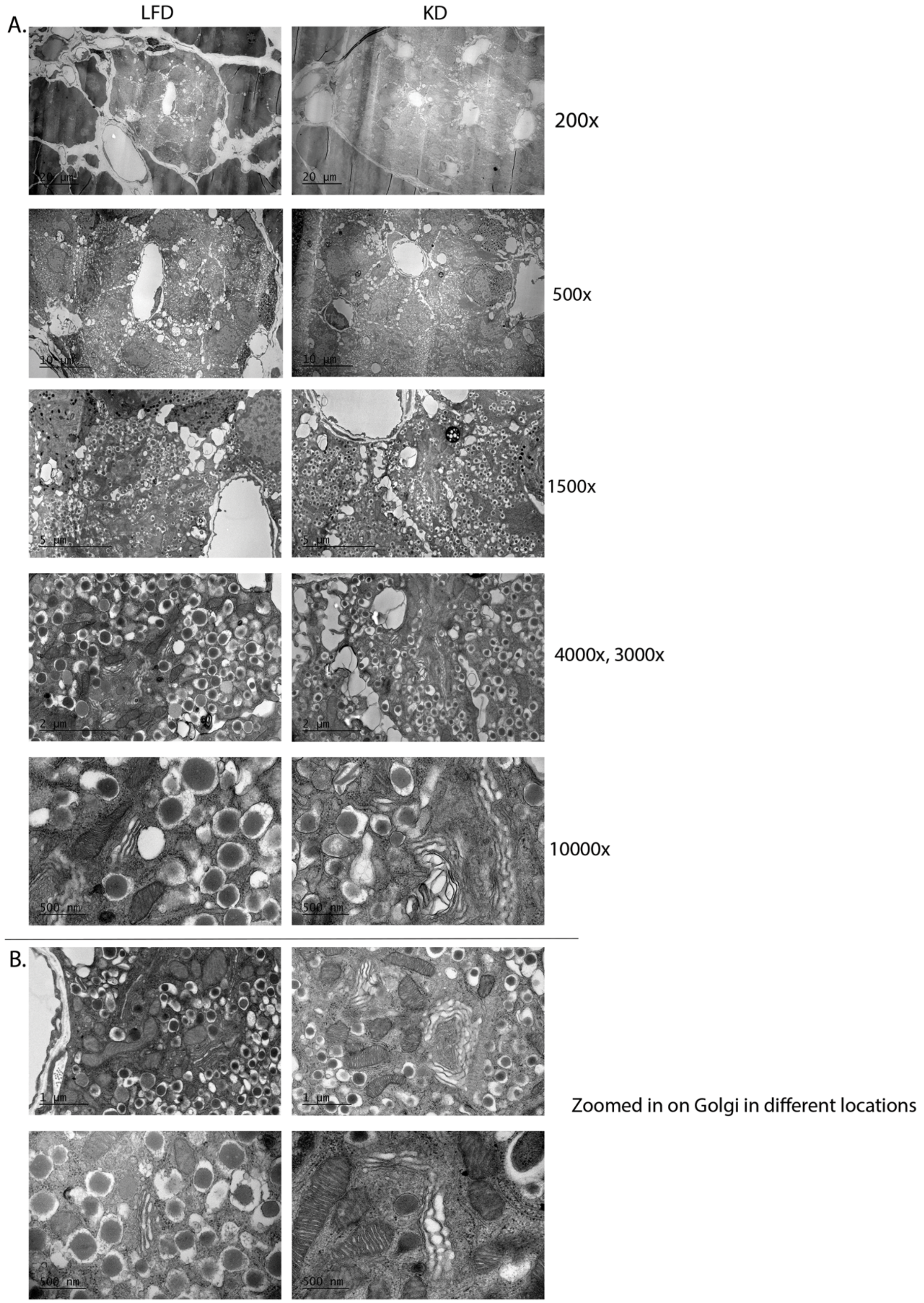

Figure S7 related to figure 7

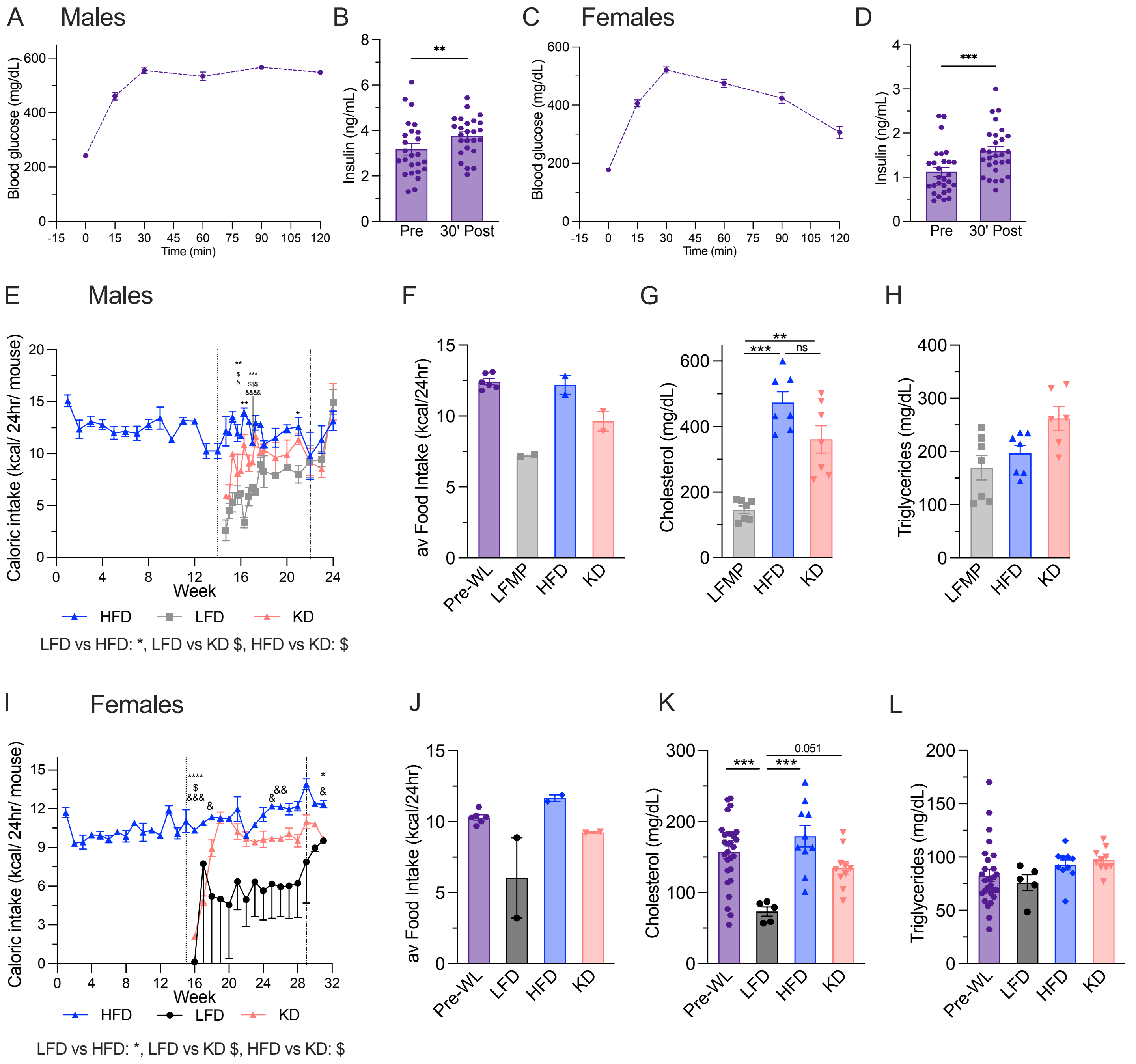
